# Benchmarking Statistical Methods for Differential Composition Analysis in Population-Scale Single-Cell Data

**DOI:** 10.64898/2026.11.04.764099

**Authors:** Weiwei Wu, Mengyu He, Yi-Juan Hu, Chang Su

## Abstract

Population-scale single-cell RNA sequencing (scRNA-seq) data capture cellular composition across cohorts ranging from several dozens to thousands of individuals, enabling the detection of cell-type composition differences between conditions. These differences reveal cellular shifts in disease pathogenesis and suggest potential targets for therapeutic intervention. However, differential composition analysis in scRNA-seq data is challenged by the unique features of cell count data, such as compositionality, prevalence of zeros, overdispersion, and confounding by technical and biological covariates. While this analysis is routinely performed, no study has yet systematically evaluated existing methods. Here, we benchmarked 12 methods for differential composition analysis using three population-scale scRNA-seq datasets with diverse and representative characteristics. Analyses of real data revealed low agreement across methods, while permutations and simulations based on the real data demonstrated excessive false discoveries for most methods. Overall, LOCOM emerged as the top-performing method that controls false discovery rate while maintaining good statistical power, followed by ANCOM-BC and linear regression with centered-log-ratio transformation. Integrating evidence from real-data analyses and data-informed permutation and simulation experiments, we provide practical guidance tailored to common real-world challenges in differential composition analysis.

## 1 Introduction

The past few years have witnessed rapid expansion of population-scale single-cell RNA sequencing (scRNA-seq) datasets, each comprising several dozens to thousands of samples across conditions in the population. **Table 1** lists more than 30 such studies. Notably, the Religious Order Study and the Rush Memory and Aging Project (ROSMAP)[1] profiled over 1.65 million nuclei from 437 brain tissue samples of Alzheimer’s disease (AD) patients and elderly individuals, and the OneK1K project collected 1.27 million peripheral blood mononuclear cells (PBMCs) from 982 healthy donors [2].

**Table 1:**
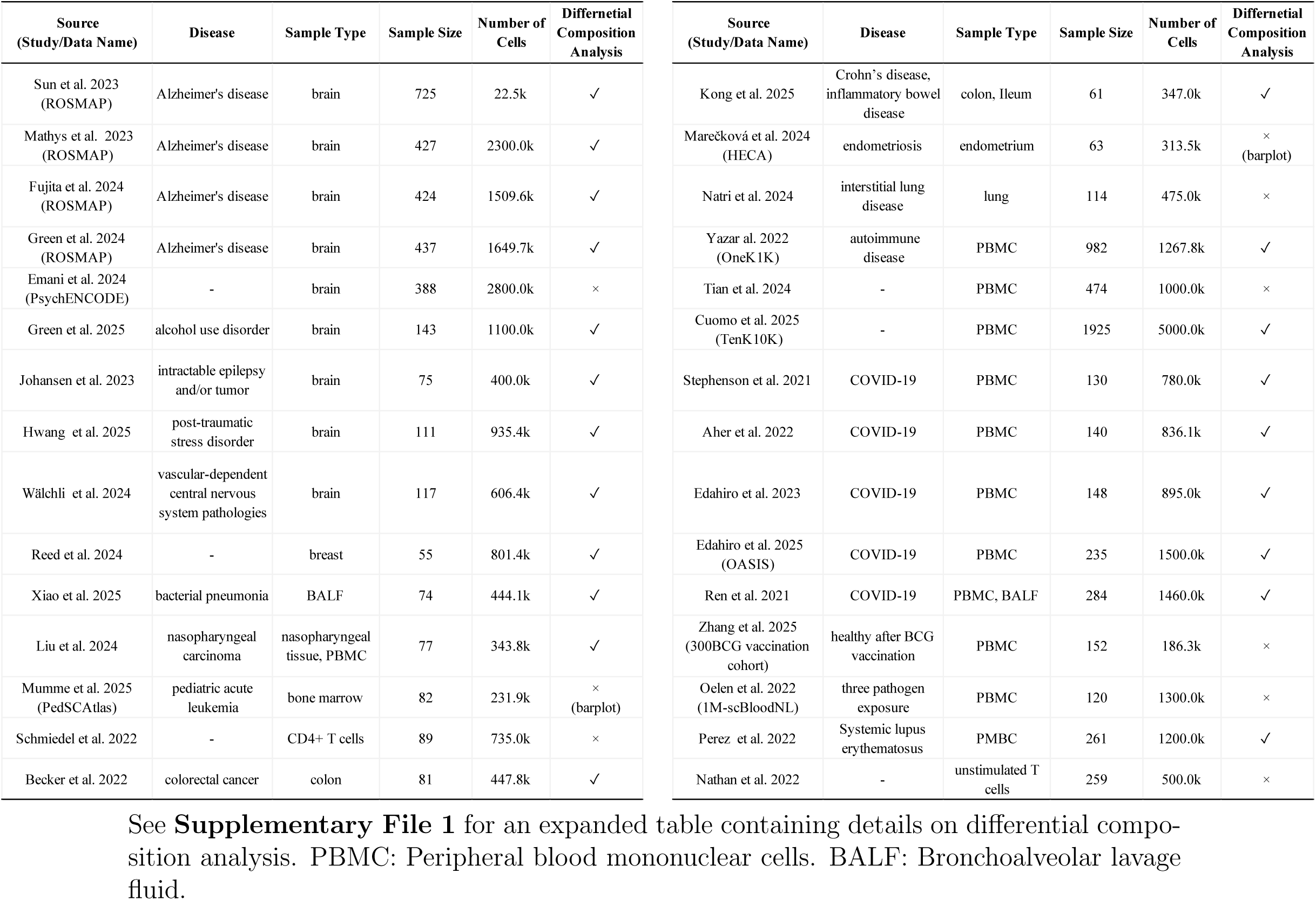
Population-scale single-cell studies.

| Source (Study/Data Name) | Disease | Sample Type | Sample Size | Number of Cells | Differential Composition Analysis | Source (Study/Data Name) | Disease | Sample Type | Sample Size | Number of Cells | Differential Composition Analysis |
|---|---|---|---|---|---|---|---|---|---|---|---|
| Sun et al. 2023 (ROSMAP) | Alzheimer's disease | brain | 725 | 22.5k | ✓ | Kong et al. 2025 | Crohn’s disease, inflammatory bowel disease | colon, Ileum | 61 | 347.0k | ✓ |
| Mathys et al. 2023 (ROSMAP) | Alzheimer's disease | brain | 427 | 2300.0k | ✓ | Marečková et al. 2024 (HECA) | endometriosis | endometrium | 63 | 313.5k | ×<br>(barplot) |
| Fujita et al. 2024 (ROSMAP) | Alzheimer's disease | brain | 424 | 1509.6k | ✓ | Natri et al. 2024 | interstitial lung disease | lung | 114 | 475.0k | × |
| Green et al. 2024 (ROSMAP) | Alzheimer's disease | brain | 437 | 1649.7k | ✓ | Yazar al. 2022 (OneK1K) | autoimmune disease | PBMC | 982 | 1267.8k | ✓ |
| Emani et al. 2024 (PsychENCODE) | - | brain | 388 | 2800.0k | × | Tian et al. 2024 | - | PBMC | 474 | 1000.0k | × |
| Green et al. 2025 | alcohol use disorder | brain | 143 | 1100.0k | ✓ | Cuomo et al. 2025 (TenK10K) | - | PBMC | 1925 | 5000.0k | ✓ |
| Johansen et al. 2023 | intractable epilepsy and/or tumor | brain | 75 | 400.0k | ✓ | Stephenson et al. 2021 | COVID-19 | PBMC | 130 | 780.0k | ✓ |
| Hwang et al. 2025 | post-traumatic stress disorder | brain | 111 | 935.4k | ✓ | Aher et al. 2022 | COVID-19 | PBMC | 140 | 836.1k | ✓ |
| Wälchli et al. 2024 | vascular-dependent central nervous system pathologies | brain | 117 | 606.4k | ✓ | Edahiro et al. 2023 | COVID-19 | PBMC | 148 | 895.0k | ✓ |
| Reed et al. 2024 | - | breast | 55 | 801.4k | ✓ | Edahiro et al. 2025 (OASIS) | COVID-19 | PBMC | 235 | 1500.0k | ✓ |
| Xiao et al. 2025 | bacterial pneumonia | BALF | 74 | 444.1k | ✓ | Ren et al. 2021 | COVID-19 | PBMC, BALF | 284 | 1460.0k | ✓ |
| Liu et al. 2024 | nasopharyngeal carcinoma | nasopharyngeal tissue, PBMC | 77 | 343.8k | ✓ | Zhang et al. 2025 (300BCG vaccination cohort) | healthy after BCG vaccination | PBMC | 152 | 186.3k | × |
| Mumme et al. 2025 (PedSCAtlas) | pediatric acute leukemia | bone marrow | 82 | 231.9k | ×<br>(barplot) | Oelen et al. 2022 (1M-scBloodNL) | three pathogen exposure | PBMC | 120 | 1300.0k | × |
| Schmiedel et al. 2022 | - | CD4+ T cells | 89 | 735.0k | × | Perez et al. 2022 | Systemic lupus erythematosus | PMBC | 261 | 1200.0k | ✓ |
| Becker et al. 2022 | colorectal cancer | colon | 81 | 447.8k | ✓ | Nathan et al. 2022 | - | unstimulated T cells | 259 | 500.0k | × |

See **Supplementary File 1** for an expanded table containing details on differential compo-sition analysis. PBMC: Peripheral blood mononuclear cells. BALF: Bronchoalveolar lavage fluid.

By capturing thousands of cells per sample, cellular composition can be systematically quantified at the population level. This presents an unprecedented opportunity to identify condition-associated cellular shift through differential composition analysis, offering insights into the underlying biological processes. Indeed, differential composition analysis has been widely performed in recent population-scale single-cell studies (**Table 1**). For instance, the ROSMAP single-cell cohort [1] identified microglia to be associated with AD-related endophenotypes. Similarly, a multi-omic blood atlas of COVID-19 revealed severity-dependent alterations across multiple immune cell populations, including neutrophils, progenitor cells, and lymphocytes [3]. **Supplementary File 1** details the differential composition analyses conducted in the studies summarized in **Table 1**, including the conditions of interest and the statistical methods used.

**Fig. 1** illustrates a general workflow of differential composition analysis. Briefly, single cells across samples are profiled by scRNA-seq, then clustered and assigned to cell types. Differential composition analysis then takes as input a cell count matrix, which records the number of cells for each cell type in each sample, to detect the difference in cell-type composition between conditions. This count matrix has several unique characteristics that pose challenges for differential composition analysis. First, the total number of cells captured in a sample (referred to as “library size”, a term borrowed from sequencing data analysis) is strongly influenced by technical factors [4] and may not reflect the true absolute number of cells in the sample. Consequently, analyses typically focus on relative abundances to mitigate the impact of varying library sizes. However, the relative abundances of cell types are compositional (constrained to sum to one within each sample), so that a change in one cell type’s abundance can induce spurious changes in others [5, 6]. This makes it challenging to identify the causal cell types whose abundances truly differ between conditions and to guard against false discoveries driven by the compositional constraint. Second, cell count data contain a large number of zeros, as rare or low-abundant cell types may be absent from a subset of samples [7]. The prevalence of zeros is known to cause analytical difficulties [8]. In addition, overdispersion refers to the greater-than-expected variability in cell counts or proportions across samples. This often arises from biological and sampling variability and can lead to underestimated variance and inflated false positives [9, 10]. Moreover, testing tens to hundreds of cell types simultaneously requires multiple-testing correction, which becomes more difficult when the cell types are correlated [11, 12]. Finally, technical and biological covariates associated with the condition of interest may confound differential composition analysis if unaccounted for. For example, differential library sizes across conditions have been shown to bias estimates of compositional differences [13], and biological covariates such as age, sex, and tissue source may similarly confound the analyses. We note that these data characteristics closely mirror those commonly encountered in microbiome sequencing data [14].

**Figure 1:**
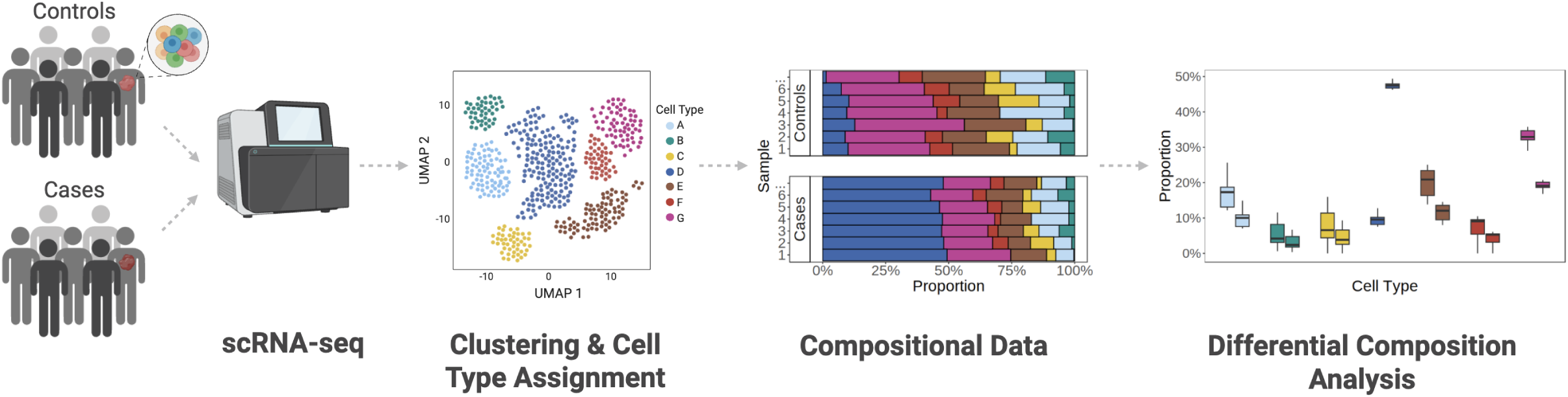
Pipeline of differential composition analysis for scRNA-seq data. Cells are clustered and annotated into cell types based on single-cell gene expression data. Relative abundances of these cell types are quantified per sample, followed by statistical testing to identify significant differences between conditions. Created with BioRender.com.

A variety of methods have been used to differential composition analysis of cell count data (**Table 2**). Some were specifically developed for scRNA-seq cell count data, such as DCATS [15], propeller [16], sccomp [17], and scCODA [18]. Others were originally developed for microbiome data, such as LOCOM [6], ANCOM-BC [19], ANCOM-BC2 [13], or bulk RNA-seq data, such as DESeq2 [20]. In addition, conventional statistical approaches have also been applied, including linear regression (LR), the Wilcoxon test, and the quasi-binomial F-test [3, 21, 22]. These methods differ substantially in how they handle compositional constraints, excessive zeros, and overdispersion. Some explicitly account for the compositional nature of the data through log-ratio transformations (LOCOM, sccomp), log-transformations with constraints (scCODA), bias correction (ANCOM-BC, ANCOM-BC2), or normalization (DE-Seq2, DCATS). In contrast, others analyze each cell type’s relative abundance independently (propeller, quasi-binomial F-test). While certain methods (e.g., LOCOM, DCATS, quasi-binomial F-test) analyze zeros as they are, many others require adding pseudo-counts. Finally, overdispersion is addressed either nonparametrically (LOCOM, Wilcoxon) or through parametric models with explicit overdispersion parameters. Many methods employ the Benjamini-Hochberg (BH) procedure [23] to adjust for multiple tests; however, this procedure assumes that the tests are independent or follow a positive-regression-dependence structure (PRDS) [24].

**Table 2:** Twelve methods for differential composition analysis evaluated in this study.

|  | **For general compositional data** |  |  | **For single cell data** |  |  |  | **For microbiome data** |  |  |  |
|---|---|---|---|---|---|---|---|---|---|---|---|
| **Methods** | **LR+CLR** | **Wilcoxon+CLR** | **Quasi-binomial F-Test** | **DCATS** | **propeller** | **sccomp<br>(sccomp-or)** | **scCODA** | **LOCOM** | **ANCOM-BC** | **ANCOM-BC2** | **DESeq2** |
| **Data** | CLR | CLR | Count | Count | Logit/arcsin sqrt | Count | Count | Count | Log | Log | Count |
| **Distribution assumption** | - | - | Binomial | Beta-binomial | - | Sum-constrained Beta-binomial | Dirichlet-multinomial | - | - | - | Negative Binomial |
| **Model** | Linear | Rank | Logistic | GLM | Linear | Bayesian | Bayesian | Logistic | Linear | Linear | GLM |
| **Addressing compositionality** | ✓ | ✓ | × | ✓ | × | ✓ | ✓ | ✓ | ✓ | ✓ | ✓ |
| **Adjusting confounders** | ✓ | × | ✓ | ✓ | ✓ | ✓ | ✓ | ✓ | ✓ | ✓ | ✓ |
| **Allowing continuous traits** | × | × | × | ✓ | × | × | ✓ | ✓ | × | ✓ | ✓ |
| **Bypassing pseudo-count** | × | × | ✓ | ✓ | × | ✓ | ✓ | ✓ | × | × | ✓ |
| **Reference** |  |  | Pal et al., 2021 | Lin et al., 2023 | Phipson et al., 2022 | Mangiola et al., 2023 | Buttner et al., 2021 | Hu et al., 2022 | Lin and Peddada, 2022 | Lin and Peddada, 2024 | Love et al., 2014 |

LR: linear regression. CLR: centered log-ratio transformation. GLM: generalized linear model. All methods were developed in R, except for scCODA, which was developed in Python.

Despite the growing number of available methods and the widespread use of differential composition analysis in population-scale single-cell studies, no systematic benchmarking has been conducted to evaluate the performance of existing methods on cell count data from scRNA-seq. While cell count data share similarity with taxonomic count data from microbiome studies, benchmarking studies in the microbiome field [5, 14] do not faithfully capture the characteristics of cell count data, nor evaluate the newer or cell-count-specific methods. This lack of evaluation leaves users with limited guidance for selecting the most appropriate approach for differential composition analysis in population-scale single-cell studies.

To address this gap, we benchmarked 12 differential composition analysis methods, based on three large-scale single-cell datasets with diverse biological conditions and technical characteristics. Our evaluation spanned real data analysis, permutation experiments, and seven simulation settings designed to reflect varying analytical challenges. Based on our findings, we provided practical recommendations for method selection and usage. We also highlighted limitations and future directions in single-cell differential composition analysis to support the continued development of analytic tools for this field.

## 2 Results

### 2.1 Differential Composition Methods

In this study, we systematically benchmarked 12 methods. These include DCATS, propeller, sccomp, sccomp with an outlier-removal step (sccomp-or), scCODA, LOCOM, ANCOM-BC, ANCOM-BC2, DESeq2, LR and Wilcoxon rank-sum tests on centered-log-ratio (CLR) transformed data (LR+CLR, Wilcoxon+CLR), and the quasi-binomial F-test. To facilitate comparison across methods, we summarized their key features in **Table 2** and provide further discussion below.

Log-ratio transformations, such as CLR and additive-log-ratio (ALR), are effective ways to mitigate the compositional constraint inherent in relative abundance data [25, 26]. LR+CLR and Wilcoxon+CLR apply the CLR transformation directly to the observed count data, whereas LOCOM and sccomp embed these transformations in the parameter space of their statistical models. In a similar spirit to the log-ratio transformation, ANCOM-BC and ANCOM-BC2 apply the log transformation to the count data followed by bias-correction, while scCODA log-transforms its underlying parameters and constrains the coefficient corresponding to a chosen reference category. DESeq2 and DCATS instead attempt to normalize read counts to approximate absolute abundances, thereby circumventing the compositional constraints of relative abundance data; however, these estimates rely on the appropriate selection of a normalization baseline or reference population [27]. As non-compositional approaches, propeller fits linear regressions to logit- or arcsine-square-root transformed proportions, while the quasi-binomial F-test fits logistic regressions to count data; these approaches treat each cell type independently and may fail to account for the compositional dependency among cell types [28].

All methods that rely on log or logit transformations of observed count data face a common issue: the presence of zero counts, which cannot be transformed. A common workaround is to add a pseudo-count to zeros or to all counts. Although this enables the transformation, previous studies have shown that pseudo-count addition can introduce systematic bias, thereby inflating error rates in differential composition analyses [6, 29]. An alternative approach is to apply log or log-ratio transformations to the model parameters while treating zeros as natural observations (LOCOM and sccomp).

In some of these methods, overdispersion is addressed by a parametric approach that extends simple binomial or Poisson distribution to quasi-binomial (quasi-binomial F-test), beta-binomial (DCATS and sccomp), Dirichlet-multinomial (scCODA), or negative-binomial distribution (DESeq2). However, it may be challenging to identify a single parametric distribution that adequately fits data from all cell types [17]. Similarly, LR, propeller, ANCOM-BC and ANCOM-BC2 are based on linear regressions, implicitly assuming (approximate) normality for the CLR, logit/arcsin-square-root, or log transformed data, which does not always hold [30]. Other methods take a nonparametric approach and may be more robust: propeller employs a variance-stabilization procedure through empirical Bayes shrinkage; LOCOM accommodates overdispersion by adopting the generalized estimating equation framework.

To account for multiple testing in high-dimensional settings, all non-Bayesian methods except for ANCOM-BC and ANCOM-BC2 employ the BH procedure to control the false discovery rate (FDR). However, the BH procedure assumes that the tests are independent or follow a positive-regression-dependence structure (PRDS), which may not be satisfied by cell types that are inter-dependent in complex ways. In contrast, ANCOM-BC and ANCOM-BC2 by default use the Holm-Bonferroni method [12], which controls the family-wise error rate (FWER) and is therefore more conservative than FDR control. For consistency and fair comparison across methods, we applied the BH correction to all non-Bayesian differential composition methods in this study. The two Bayesian approaches, scCODA and sccomp, assess significance by borrowing information across cell types and reporting FDR-adjusted results according to their respective criteria.

Finally, some methods incorporate offsets or normalization procedures to mitigate batch effects and differential library sizes across conditions (e.g., ANCOM-BC2). Among the methods evaluated, all support adjustment for covariates, except for Wilcoxon+CLR. By default, all methods are designed to test differential composition across conditions, which is a binary trait. LR+CLR, DCATS, scCODA, LOCOM, ANCOM-BC2, and DESeq2 can also accommodate continuous traits of interest. Details on the implementation of each method can be found in the “Method Implementation” section of Methods.

### 2.2 Benchmarking Framework

**Fig. 2** illustrates our benchmarking framework. We collected three real datasets with diverse characteristics to evaluate method performance in realistic settings. The framework consists of three modules. In the first module, we assessed the consistency of results across 12 methods on real data, which revealed their reliability in practical settings. In the second module, we created null datasets, where all cell types are independent of the trait of interest, through permuting the trait values in the real datasets, to assess the methods’ control of false-positive error rates (FDR and type I error rate) under a global null setting. This was followed by the third module, where we simulated alternative datasets that incorporated differential compositions to evaluate the methods’ sensitivity. In this module, we designed seven settings that reflect analytical challenges in single-cell studies, such as differential library sizes, correlations among cell types, the presence of confounders or continuous traits, and small sample sizes. We also carefully selected compositional data simulators to best capture the real-data features. Finally, we summarized the performance of different methods using a ranking-based approach.

**Figure 2:**
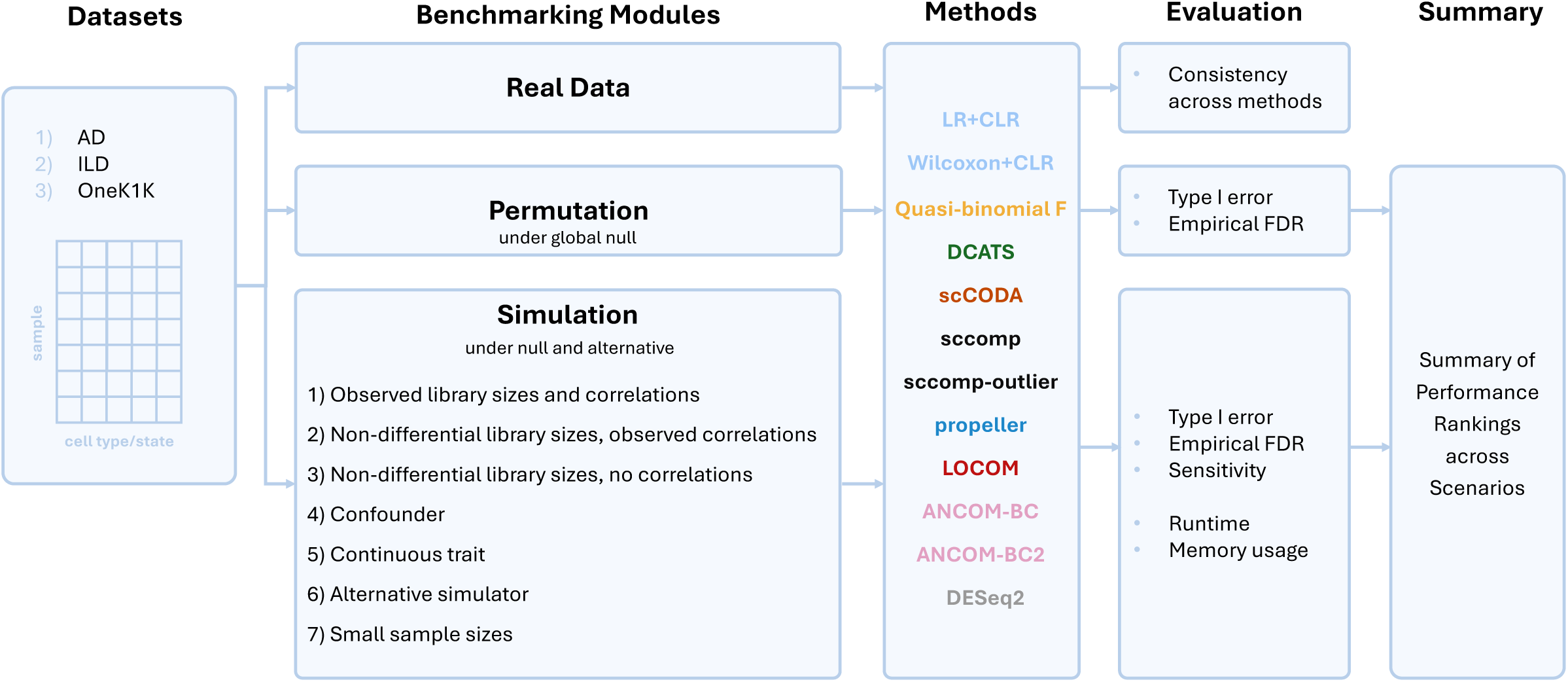
An overview of the benchmarking framework. Using three real datasets (AD [1], ILD [31], OneK1K [2]) and a three-module benchmarking pipeline, we evaluated the performance of 12 methods across multiple settings and metrics, and summarized the results using a ranking-based approach (see “Benchmarking Metrics” section of Methods).

### 2.3 Real Datasets

**Table 3** lists three population-scale scRNA-seq datasets used in this study, with predefined cell type annotations. These datasets represent a variety of tissue types, sample sizes, and data features, including (1) the interstitial lung disease (ILD) dataset [31] on lung tissue samples from 95 ILD patients and controls, (2) the Alzheimer’s disease (AD) dataset, consisting of postmortem brain tissue samples from 437 AD patients and controls from the ROSMAP study [1], and (3) the OneK1K dataset [2] on peripheral blood mononuclear cell (PBMC) from 939 healthy participants. For the AD data, AD status was used as the trait of interest, and APOE *ɛ*4, study, postmortem interval, and sex were included as covariates. For the ILD data, lung disease status was used as the trait of interest, and age and sample source were included as covariates. Since the OneK1K dataset consists only of healthy individuals, we used the sex variable as the trait and adjusted age as the covariate. The association statistics between covariates and the traits of interest across three datasets are presented in **Supplementary Table. S2** The details of data preprocessing are presented in Section 4.1.

**Table 3:** Three population-scale scRNA-seq datasets used in this study.

| **Dataset** | **Tissue type** | **Trait** | **Number of samples** | **Cell group** | **Number of cell groups** | **Total number of cells** |
|---|---|---|---|---|---|---|
| ILD | Lung | Disease status | 95 | Cell type | 26 | 442,532 |
|  |  |  |  | Cell state | 43 | 442,532 |
| AD | Brain | Disease status | 437 | Cell type | 13 | 1,603,334 |
|  |  |  |  | Cell state | 86 | 1,592,028 |
| OneK1K | PBMC | Sex | 939 | Cell state | 26 | 1,219,020 |

AD: Alzheimer’s disease. ILD: Interstitial lung disease. PBMC: Peripheral blood mononuclear cells. The data have been preprocessed according to section “Real Data Preprocessing” of Methods.

The AD and ILD datasets have annotations at both cell type (CT) and cell state (CS) levels, with cell state being a finer functional unit within a cell type [32]. We refer to both levels as the “cell group”. AD was annotated for 13 CTs and 86 CSs, and ILD for 26 CTs and 43 CSs. In contrast, the OneK1K dataset has only CS annotations available, comprising 26 CSs. Hereafter, we treat each dataset–cell group combination in Table 3 as a distinct dataset, yielding five datasets in total. The data characteristics are displayed in **Fig. 3**, which shows substantial variability across the five datasets. First, the cell group abundance data have many zeros (**Fig. 3a**). In the AD dataset, 14.28% of CTs and 20.43% of CSs contain more than 50% of zero observations across all samples. The ILD dataset has fewer zeros than the AD and OneK1K datasets. Second, the library sizes are highly variable across samples and datasets (**Fig. 3b**). OneK1K has the smallest library sizes overall, averaging around 1,000. The ILD dataset exhibits notable differences in library sizes between cases and controls. Third, the cell group abundances can be highly correlated (**Fig. 3c**). Higher correlations between cell groups were observed in ILD at both the CT and CS levels, as well as in AD at the CS level, compared to OneK1K CS and AD CT. Fourth, the cell abundance data exhibit varying degrees of overdispersion across datasets (**Fig. 3d**). More detailed results are provided in **Supplementary Fig. S1-S3**.

**Figure 3:**
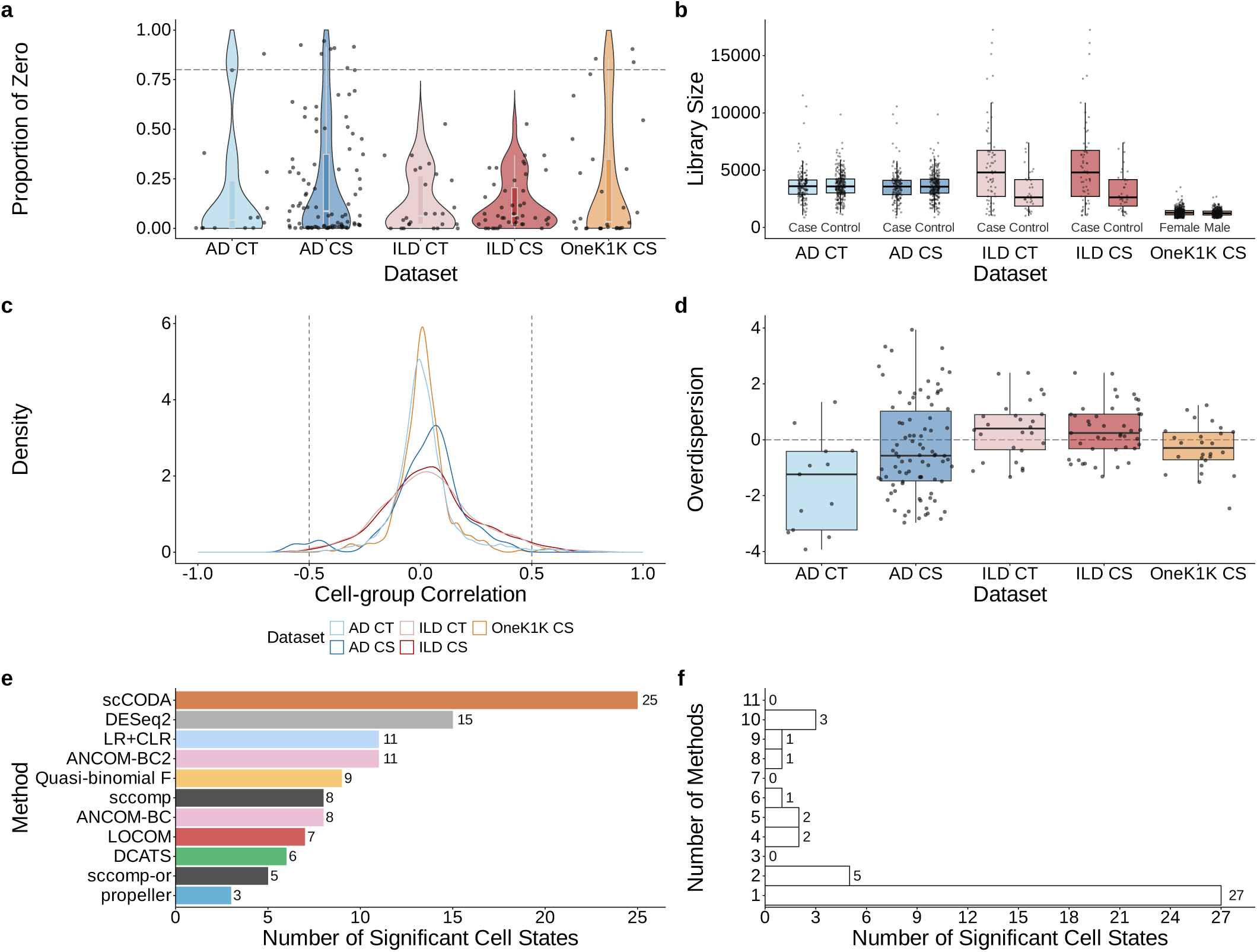
Data characteristics of the five real datasets (before data preprocessing) (**a**-**d**) and analysis results of the AD CS dataset (**e**-**f**): **a**. Proportion of zero by cell group (the horizontal dashed line at 0.8 denotes the threshold used in preprocessing to exclude cell groups with more than 80% zero proportions from downstream differential composition analyses); **b**. Library size by sample group; **c**. Pearson’s correlation between cell groups (vertical dashed lines at cell-group correlations of *±*0.5), using the CLR-transformed data and zero imputation [49]; **d**. Overdispersion by cell group (horizontal dashed line at 0, indicating no overdispersion), estimated by the overdispersion parameter from a Poisson GLM that adjusts for the trait of interest, fitted with glm() and dispersiontest() in R; **e**. Number of cell states detected by different methods in the AD CS dataset; **f**. Number of detected cell states (in AD CS) shared across varying numbers of methods. Results for the other datasets are provided in Supplementary Fig. S4.

We applied 11 methods to analyze these datasets except for Wilcoxon+CLR due to the presence of covariates. We detected cell groups whose composition differed between case and control samples based on the prespecified FDR significance threshold of 0.01. **Fig. 3e-f** show low agreement in detected cell states across 11 methods in the AD CS dataset. In particular, there are no cell states detected by all 11 methods, and only three detected cell states were shared by 10 methods and 27 were unique to one method. This low agreement was also seen on other datasets (**Supplementary Fig. S4**). The phenomenon raises concerns about the reliability of differential composition results reported in the literature. We hypothesize that this is caused by the inability of some existing methods to properly address common challenges in single-cell data, leading to inadequate control of false-positive findings. These observations motivated the permutation and simulation modules presented below.

### 2.4 Null Datasets via Permutation

We first evaluated false-positive control of different methods under the global null hypothesis. We considered five scenarios, one for each dataset-cell group combination in **Table 3**. For each scenario, we generated 1,000 global null replicates via permutation, i.e., randomly shuffling the trait of interest so that none of the cell groups is associated with the trait. For each method, the FDR with a 95% confidence interval was assessed at a nominal level of 10%; the type I error rates by cell group were assessed at the nominal 0.05 level (“Benchmarking Metrics” of Methods).

**Fig. 4a** displays the FDR results for all methods across the five scenarios. LOCOM, DCATS, propeller, LR+CLR, and Wilcoxon+CLR demonstrated proper FDR control. Among these, DCATS, propeller, LR+CLR, and Wilcoxon+CLR showed slight deflation (i.e., empirical FDRs are smaller than the lower bound of the confidence interval) in some scenarios. The quasi-binomial F-test exhibited mild inflation in the ILD scenarios at both the CT and CS levels. Sccomp and its outlier removal version showed inconsistent performance across scenarios, with substantial inflation in the ILD two scenarios, while being overly conservative in other scenarios. ANCOM-BC failed to control FDR across all scenarios, with a median FDR of 0.18, which is almost twice the nominal level that it claims to maintain. In contrast, its successor, ANCOM-BC2, generally maintained FDR within the upper bound of the confidence interval, except for a much elevated FDR (0.66) in the AD CS scenario. sc-CODA and DESeq2 exhibited substantial inflation in permutation experiments across all scenarios. Consistent with our findings, the sccomp paper [17] also reported undesirable false-positive control by scCODA. DESeq2 may have failed because it uses the negative binomial distribution, which does not adequately capture the overdispersion of cell type counts.

**Figure 4:**
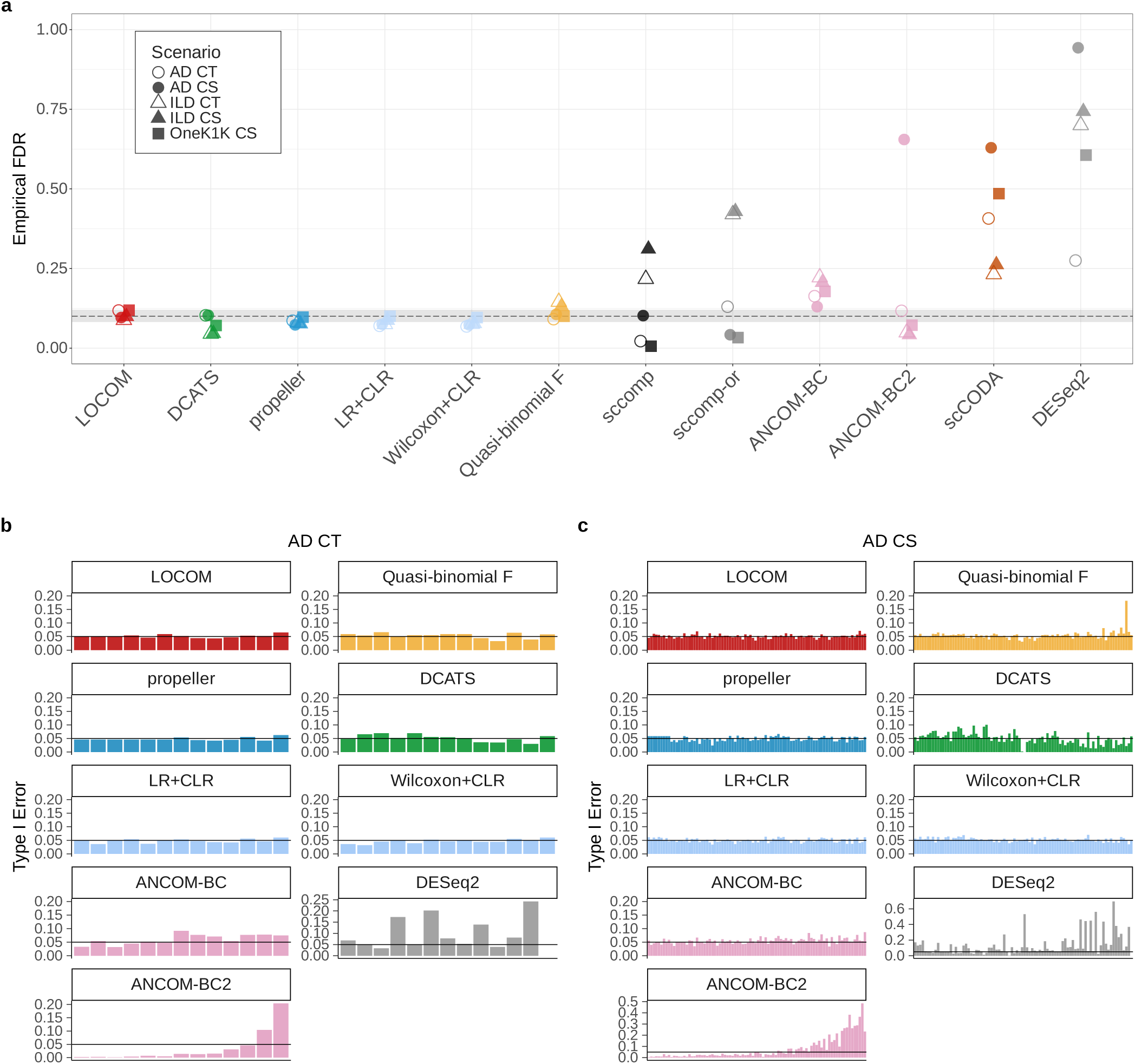
Empirical FDR in analysis of permuted (global null) replicates of all five scenarios (**a**), and type I error rates by cell group in analysis of permuted replicates of the AD CT scenario (**b**) and the AD CS scenario (**c**), respectively. The type I error rates for other scenarios are presented in Supplementary Fig. S5. In **a**, the dashed black line represents the nominal 10% level, and the gray shaded band indicates the 95% confidence interval. In **b** and **c**, the x-axes denote cell types and cell states in ascending order of the proportion of zero, respectively; the corresponding cell type/state names are provided in Supplementary Fig. S1. The black lines mark the nominal 0.05 level.

While the empirical FDR summarizes false-positive control over all cell groups, the degree of control may vary across individual cell groups. To investigate this, we further computed the type I error rate for each cell group (**Fig.4b-c**, **Supplementary Fig. S5**). An interesting example is ANCOM-BC2, which generally maintained FDR yet showed unique inflation patterns in type I errors. Its type I error rates appeared to be associated with the proportion of zeros in each cell group, generally showing deflation in cell groups with a higher proportion of zeros (e.g., Oli.3 in AD CS) and inflation in those with fewer zeros (e.g., NK cells in AD CS). This trend was consistent across all five scenarios. In contrast, other methods failed to control type I error in distinct ways. DESeq2 showed high inflation at some cell groups with no clear dependence on abundance. The quasi-binomial F-test effectively controlled type I error rates in the AD CT and OneK1K scenarios but failed in some rare cell groups in the other scenarios. Type I error rates of ANCOM-BC showed a slight and persistent inflation above the nominal level, especially for the two ILD scenarios. DCATS exhibited both inflation and deflation across different cell groups with no clear dependence on abundance. Notably, LOCOM, propeller, LR+CLR, and Wilcoxon+CLR achieved proper type I error control across cell groups and scenarios. scCODA and sccomp were excluded from the type I error evaluation as their Bayesian frameworks are not compatible with frequentist type I errors.

In summary, LOCOM, propeller, LR+CLR, and Wilcoxon+CLR demonstrated strong control of both FDR and type I error rates under a global null hypothesis. Quasi-binomial F test, sccomp, sccomp-or, and ANCOM-BC showed moderate performance, with acceptable FDR and type I error control in some scenarios but apparent inflation in others. scCODA and DESeq2 were excluded from subsequent simulation experiments due to their persistently high FDR across all scenarios.

### 2.5 Alternative Datasets via Controlled Simulation

The null datasets described above evaluate false-positive control in isolation, but in practice, datasets contain a mix of truly differential and non-differential cell types. To evaluate both sensitivity and false-positive control simultaneously, we designed a simulation module that introduces compositional differences under the alternative hypothesis.

#### 2.5.1 Simulator selection

We began by identifying a simulator that could faithfully reproduce the features of cell count data from scRNA-seq. Four data simulators were considered: the Dirichlet multinomial (DM) model, the sccomp simulator, MIDASim [33], and sparseDOSSA [34]. The DM model is widely used for simulating compositional count data [33]; the sccomp simulator is a built-in feature of the R package of sccomp; and both MIDASim and sparseDOSSA were originally developed for simulating microbiome data. Note that currently available single-cell simulators are designed for simulating gene expression data rather than cell count data [35–37] and are thus not considered.

We evaluated each simulator based on its ability to recapitulate sample-level and cell-group–level characteristics observed in real data (see “Simulator Evaluation” section of Methods). We found that MIDASim consistently generated simulated data that most closely resembled observed data. MIDASim, sccomp, and sparseDOSSA generally captured the centroid and dispersion structure of the observed samples while DM model simulated under-dispersed samples, as seen in principal coordinates analysis (PCoA) (**Supplementary Fig. S6**) and uniform manifold approximation and projection (UMAP) plots (**Supplementary Fig. S7**). This is also demonstrated by the test results from Permutational multivariate analysis of variance (PERMANOVA) (**Supplementary Table. S1**). Across cell groups, MIDASim best reproduced the proportion of zero, abundance, and overdispersion of the cell groups in five real datasets (**Supplementary Fig. S8**), while data simulated by sccomp, sparseDOSSA, and DM may not represent the observed proportion of zero or overdispersion. In addition, in terms of functionality, sccomp does not support specifying composition differences in simulations. Therefore, we selected MIDASim as the primary simulator for this study, and used sparseDOSSA as an alternative for validation, given its slower runtime and less flexible library size specification.

#### 2.5.2 Simulating differential compositions

For each of the five data-cell group scenario, we considered seven simulation settings, and generated 1,000 simulation replicates per setting and per scenario. To evaluate sensitivity and false-positive control, we selected 20% of cell groups as causal groups. For each causal group, we generated an effect size by drawing an effect magnitude from a Beta(2,2) distribution and then assigning a positive or negative direction with probabilities 0.8 and 0.2, respectively. For the AD and OneK1K scenarios, the simulated effect sizes were scaled by factors of 0.25, 0.50, 0.75, and 1. For the ILD scenarios, these scaling factors were multiplied by three to achieve sufficient power. Details of the simulation module can be found in “Simulation Details” in Methods.

#### 2.5.3 Method performance under the default simulation setting

We first simulated the default setting using library sizes and correlations among cell groups as observed in real data. The results of FDR and sensitivity are presented in **Fig. 5** and **Supplementary Fig. S17**. Note that, to facilitate fair comparisons across methods with different empirical FDR, sensitivity was calibrated at a nominal FDR level of 0.1 (see details in “Benchmarking Metrics” section of Methods), and the original sensitivity was presented in **Supplementary Fig. S9**.

**Figure 5:**
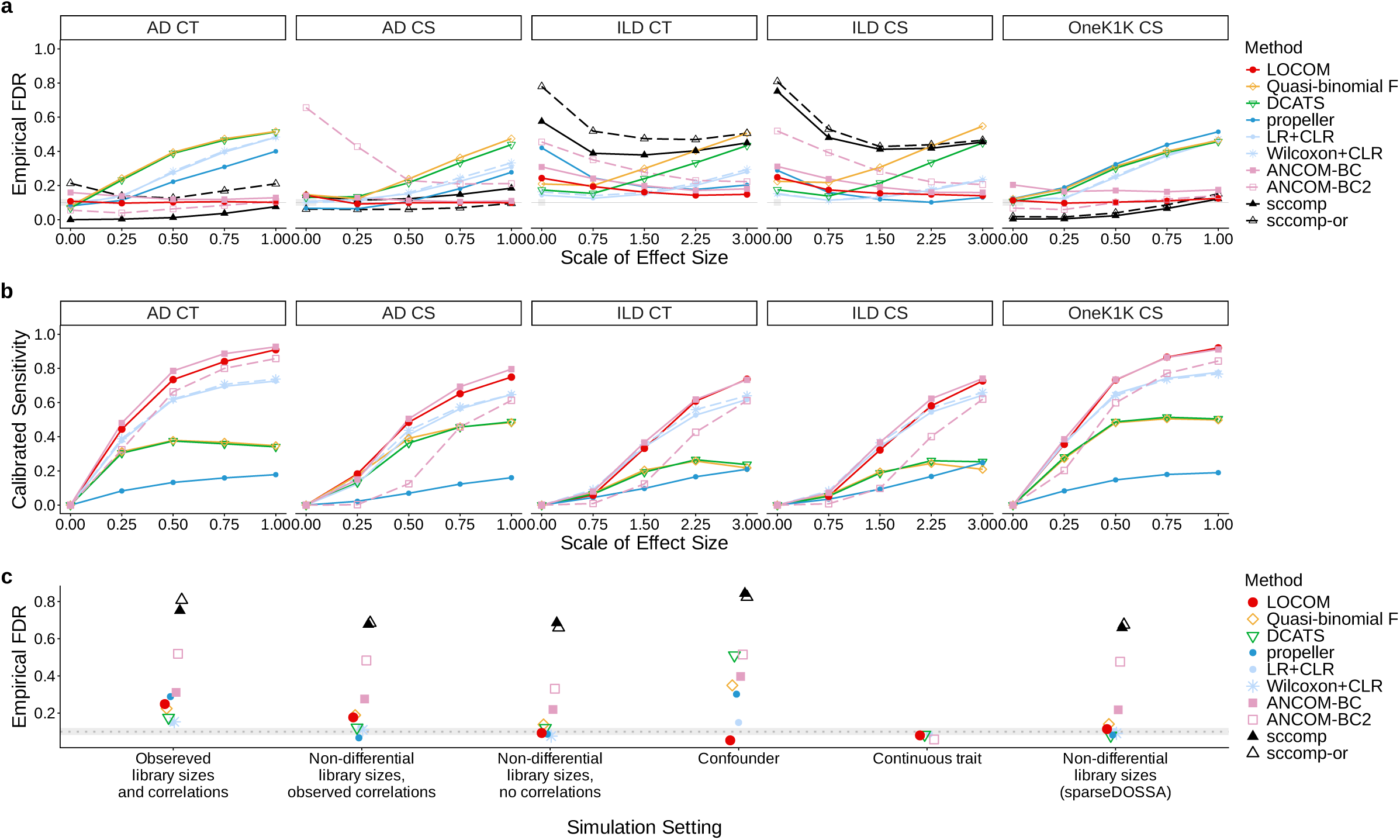
Empirical FDR and calibrated sensitivity in analysis of simulated replicates of five real datasets in the default setting (**a**-**b**), and empirical FDR in analysis of simulated null replicates of the ILD CS dataset (**c**) across six different settings. In **a**, the gray line represents the nominal 10% level and the gray shaded band indicates the 95% confidence interval. Details for evaluating calibrated sensitivity in **b** are provided in the “Benchmarking Metrics” section of Methods.

Overall, LOCOM showed the strongest FDR control, maintaining FDR close to the nominal level across nearly all scenarios, with only slight inflation in the ILD scenarios (**Fig. 5a**). ANCOM-BC also presented relatively good FDR control. FDR control was more challenging for most other methods, particularly in the ILD scenarios. LR+CLR, Wilcoxon+CLR, DCATS, and the quasi-binomial F-test exhibited FDR inflation, with the degree of inflation generally increasing as the effect size increased. In terms of calibrated sensitivity, ANCOM-BC, LOCOM, Wilcoxon+CLR, and LR+CLR achieved high sensitivity across all scenarios (**Fig. 5b**). In contrast, propeller, DCATS, and the quasi-binomial F-test showed low calibrated sensitivity in most scenarios.

We further compared the type I error rates under the global null hypothesis (i.e., scaling factor = 0) with the permutation module and found them to be generally consistent (**Fig. 5a** and **Supplementary Fig. S10**), which suggests the concordance of permutation and simulation modules and supports the validity of the evaluation framework.

#### 2.5.4 Impact of differential library sizes

Given that all methods had undesirable FDR control in the ILD scenarios (**Fig. 5a**), and that the ILD datasets had significant differences in library sizes between case and control groups (p-value *<* 0.01, Wilcoxon rank-sum test; **Fig. 3b**), we speculated that the reduced performance of differential composition methods might stem from their inability to account for differential library sizes between conditions. To test this hypothesis, we simulated a setting with non-differential library sizes while preserving all other characteristics observed in real data (**Supplementary Fig. S11**). This allowed direct comparison with the default setting (**Fig. 5**) to isolate the impact of differential library size. Indeed, the FDRs for most methods are closer to the nominal level in the ILD CS and CT scenarios (**Fig. 5c** and **Supplementary Fig. S11**), particularly when differential signals were weak. However, despite the removal of differential library sizes, the FDR of sccomp-or, sccomp, ANCOM-BC2, ANCOM-BC, DCATS, and quasi-binomial F test remained notably inflated for ILD scenarios, suggesting that additional factors may underlie their inflated false-positives.

#### 2.5.5 Impact of cell-group correlations

Considering the correlations across cell groups (**Fig. 3c**, **Supplementary Fig. S2**), which may invalidate the assumption of multiple testing correction methods [23, 24], we built upon the simulation setting with non-differential library sizes and further removed the correlations between cell groups by setting them to zero. Comparing the two settings (**Supplementary Fig. S11-S12**), we found that the FDR of the ANCOM-BC, ANCOM-BC2, quasi-binomial F test, and LOCOM decreased slightly after removing correlations, particularly when differential signals were weak. After removing both differential library sizes and cell-group correlations, only LOCOM controlled FDR across all scenarios and scales of effect size. Little change in FDR was observed in the OneK1K CS scenario, which had low correlations in the real data. No apparent changes in calibrated sensitivity were observed across scenarios after removing cell-group correlations.

#### 2.5.6 Simulations with confounders

In addition to the trait of interest, cell composition can be affected by other covariates, which may confound the results of differential composition analysis if not accounted for. However, the ability of existing methods to adjust for confounders has not been systematically evaluated. To address this, we designed a simulation setting that included a confounder correlated with the trait of interest (see “Simulation Details” section of Methods). We evaluated all methods except for the Wilcoxon+CLR, which does not support covariate adjustment. LOCOM continued to perform well across all scenarios in terms of FDR control (**Supplementary Fig. S13**), with light deflation in the ILD scenarios. By comparison, DCATS, ANCOM-BC2, ANCOM-BC, and sccomp-or showed worse FDR control compared to the default setting without a confounder (**Fig. 5**). Notably, DCATS was highly sensitive to confounding, with performance deteriorating substantially across all scenarios and FDR approaching 1 in the AD CS scenario. Interestingly, quasi-binomial F test and LR+CLR exhibited improved FDR control in the ILD CT and CS scenarios. For all methods, calibrated sensitivity was generally lower in the presence of a confounder.

#### 2.5.7 Simulations with continuous traits

Although most methods target binary traits, some diseases are more appropriately characterized by a continuous variable rather than a binary diagnosis. Nevertheless, the performance of differential composition methods for testing associations with continuous traits has not been benchmarked. To bridge this gap, we introduced a simulation setting that replaced the binary trait in the default setting with a continuous trait drawn from a standard normal distribution (see “Simulation Details” section of Methods). We evaluated LOCOM, DCATS, LR+CLR, and ANCOM-BC2, which are the only four methods among those considered that explicitly support testing associations between cell composition and continuous traits. Overall, results were consistent with the default binary setting (**Supplementary Fig. S14**, **Fig. 5**). ANCOM-BC2 continued to show highly inflated FDR in the AD CS scenario, while DCATS and LR+CLR exhibited increasing FDR inflation across all scenarios as the effect-size scale increased. In contrast, LOCOM performed the best in this setting, consistently controlling FDR and maintaining high calibrated sensitivity across five scenarios.

#### 2.5.8 Simulations with an alternative simulator: sparseDOSSA

To ensure the robustness of our benchmarking results across different simulators, we also employed sparseDOSSA as an alternative simulator. Since sparseDOSSA only supports simulations with non-differential library sizes, we used it to generate results for the setting of non-differential library sizes and observed correlations.

Trends in FDR and calibrated sensitivity (**Supplementary Fig. S15**) were consistent with those in the MIDASim-based setting (**Supplementary Fig. S11**). No major differences were observed, except for propeller, which showed improved FDR control using sparseDOSSA. These results support the generalizability of our benchmarking conclusions across different data simulators.

#### 2.5.9 Simulations with small sample sizes

Although this benchmarking study focuses on differential composition analysis for emerging population-scale data, many current single-cell studies still involve smaller sample sizes. To evaluate method performance in these cases, we simulated a small sample size setting with sample sizes of 10, 30, 50, and 70 (see “Simulation Details” in Methods).

Overall, LOCOM, LR+CLR, and Wilcoxon+CLR were the top-performing methods in this setting (**Supplementary Fig. S16**). Notably, when the sample size was 10, all methods that maintained desirable FDR control exhibited extremely low sensitivity. This implies that a sample size of 10 may be insufficient for powerful differential composition analysis, despite its frequent use in real-world studies.

### 2.6 Benchmarking Summary

To summarize benchmarking results, we proposed a rank-based approach to evaluate the relative performance of different methods by FDR and sensitivity, respectively, with smaller ranks indicating better performance. For FDR, methods were scored and ranked according to the magnitude of empirical FDR inflation above the nominal level. For sensitivity, methods were ranked based on calibrated sensitivity. Within each simulation setting, ranks were averaged across alternative datasets with varying effect sizes and combined with rankings from the null datasets. These combined ranks were summed across all scenarios (and sample sizes) to generate the final overall rankings. The rankings for each setting in the permutation and simulation modules are presented in **Table 4**, with full results available in **Supplementary File 2**.

**Table 4:**
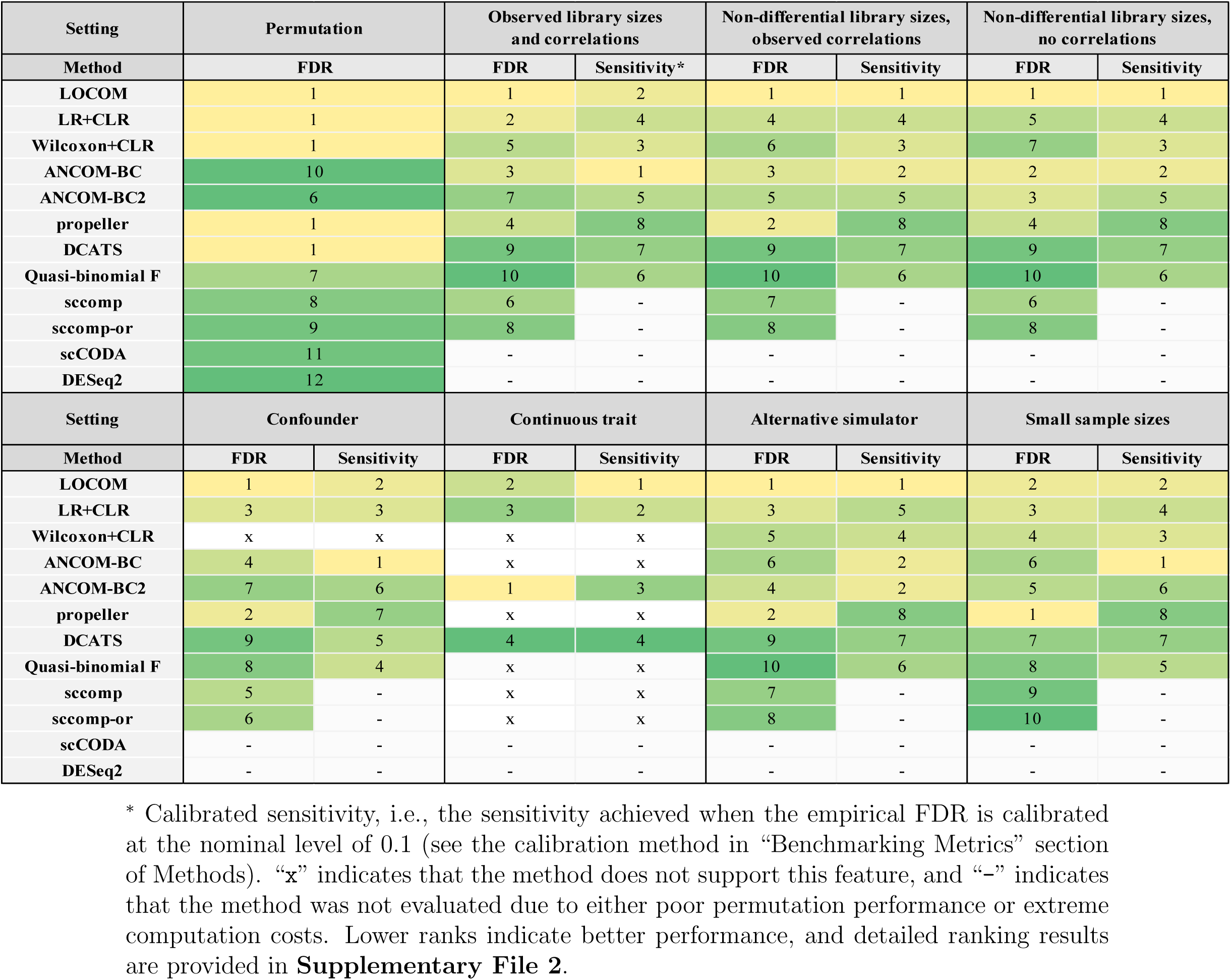
Method rankings in permutation and simulation settings.

| **Setting** | **Permutation** |  | **Observed library sizes and correlations** |  | **Non-differential library sizes, observed correlations** |  | **Non-differential library sizes, no correlations** |  |
|---|---|---|---|---|---|---|---|---|
| **Method** | **FDR** |  | **FDR** | **Sensitivity\*** | **FDR** | **Sensitivity** | **FDR** | **Sensitivity** |
| **LOCOM** | 1 |  | 1 | 2 | 1 | 1 | 1 | 1 |
| **LR+CLR** | 1 |  | 2 | 4 | 4 | 4 | 5 | 4 |
| **Wilcoxon+CLR** | 1 |  | 5 | 3 | 6 | 3 | 7 | 3 |
| **ANCOM-BC** | 10 |  | 3 | 1 | 3 | 2 | 2 | 2 |
| **ANCOM-BC2** | 6 |  | 7 | 5 | 5 | 5 | 3 | 5 |
| **propeller** | 1 |  | 4 | 8 | 2 | 8 | 4 | 8 |
| **DCATS** | 1 |  | 9 | 7 | 9 | 7 | 9 | 7 |
| **Quasi-binomial F** | 7 |  | 10 | 6 | 10 | 6 | 10 | 6 |
| **sccomp** | 8 |  | 6 | - | 7 | - | 6 | - |
| **sccomp-or** | 9 |  | 8 | - | 8 | - | 8 | - |
| **scCODA** | 11 |  | - | - | - | - | - | - |
| **DESeq2** | 12 |  | - | - | - | - | - | - |
| **Setting** | **Confounder** |  | **Continuous trait** |  | **Alternative simulator** |  | **Small sample sizes** |  |
| **Method** | **FDR** | **Sensitivity** | **FDR** | **Sensitivity** | **FDR** | **Sensitivity** | **FDR** | **Sensitivity** |
| **LOCOM** | 1 | 2 | 2 | 1 | 1 | 1 | 2 | 2 |
| **LR+CLR** | 3 | 3 | 3 | 2 | 3 | 5 | 3 | 4 |
| **Wilcoxon+CLR** | x | x | x | x | 5 | 4 | 4 | 3 |
| **ANCOM-BC** | 4 | 1 | x | x | 6 | 2 | 6 | 1 |
| **ANCOM-BC2** | 7 | 6 | 1 | 3 | 4 | 2 | 5 | 6 |
| **propeller** | 2 | 7 | x | x | 2 | 8 | 1 | 8 |
| **DCATS** | 9 | 5 | 4 | 4 | 9 | 7 | 7 | 7 |
| **Quasi-binomial F** | 8 | 4 | x | x | 10 | 6 | 8 | 5 |
| **sccomp** | 5 | - | x | x | 7 | - | 9 | - |
| **sccomp-or** | 6 | - | x | x | 8 | - | 10 | - |
| **scCODA** | - | - | - | - | - | - | - | - |
| **DESeq2** | - | - | - | - | - | - | - | - |

<sup>\*</sup> Calibrated sensitivity, i.e., the sensitivity achieved when the empirical FDR is calibrated at the nominal level of 0.1 (see the calibration method in “Benchmarking Metrics” section of Methods). “**x**” indicates that the method does not support this feature, and “–” indicates that the method was not evaluated due to either poor permutation performance or extreme computation costs. Lower ranks indicate better performance, and detailed ranking results are provided in **Supplementary File 2**.

In the permutation module, LOCOM, DCATS, LR+CLR, Wilcoxon+CLR, and propeller shared the top rank (**Table 4**). In the simulation module, under the default setting, LOCOM, ANCOM-BC, and the LR+CLR showed the strongest FDR control while maintaining relatively high sensitivity. The rankings for FDR control shifted slightly for several methods after differential library sizes and cell-group correlations were removed, but the overall trends remained similar.

LOCOM and ANCOM-BC ranked best in the confounder setting, while LOCOM and ANCOMBC2 were the top two methods in the continuous trait setting (**Table 4**). In the replication of the non-differential library sizes setting using sparseDOSSA, rankings were generally consistent with those obtained using MIDASim. Lastly, LOCOM, LR+CLR, and Wilcoxon+CLR performed well in the small-sample-size setting, ranking favorably for both FDR control and sensitivity.

Finally, we evaluated the runtime and memory usage of the 10 methods included in the simulation module for both binary and continuous traits, using 50 simulated alternative replicates of the AD CS dataset under the default setting with observed effect sizes.

For the binary trait, propeller, Wilcoxon+CLR, LR+CLR, and the quasi-binomial F-test performed best in terms of runtime. DCATS and LOCOM also ran efficiently, with average execution times under 20 seconds (**Fig. 6a**). ANCOM-BC and ANCOM-BC2 required over 3 minutes on average. sccomp and sccomp-or were the most computationally intensive, with average execution times exceeding 25 minutes, due to their underlying Bayesian framework. In terms of memory usage, propeller was the most memory-efficient method (**Fig. 6b**). In contrast, sccomp and sccomp-or showed substantial memory usage, averaging 5.50 GiB and 13.19 GiB, respectively.

**Figure 6:**
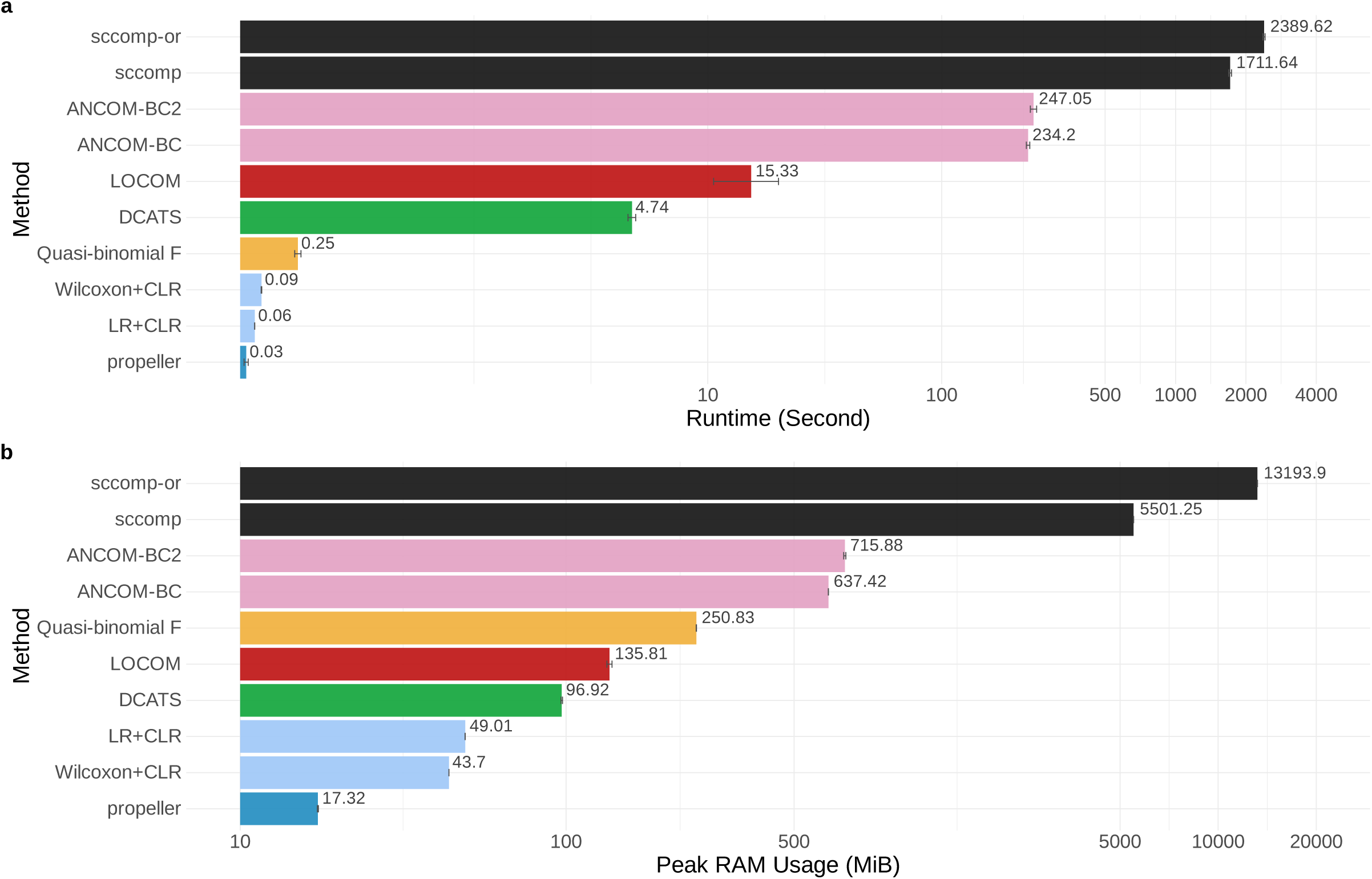
Runtime and memory usage in analysis of simulated alternative replicates of the AD CS dataset with a binary trait (**a**-**b**). The length of each bar represents the mean, and the error bars denote 1 standard deviation across replicates. Values are shown on a symmetric log scale, where a log transformation is applied for values greater than 1. All methods were run on a single core of an Intel(R) Xeon(R) Gold 6326 CPU @ 2.90GHz. Peak memory usage was monitored using the R function gc().

For the continuous trait, all three methods capable of handling continuous traits ran efficiently, with average execution times under 15 seconds (**Supplementary Fig. S18**). However, ANCOM-BC2 exhibited much higher average memory usage compared to DCATS, LOCOM, and LR+CLR.

## 3 Discussion

We systematically benchmarked 12 differential composition analysis methods using real data, permutations, and simulations, based on multiple population-scale scRNA-seq datasets with diverse characteristics.

Our findings underscore the importance of carefully selecting appropriate methods for differential composition analysis in single-cell studies. Real data analysis revealed low consistency among findings of different methods, and half of the methods exhibited inflated FDR and type I error rates in permutations. Several widely used tools also demonstrated limited performance. For example, DESeq2 and scCODA showed severely inflated FDRs. LR and the Wilcoxon rank-sum test with CLR transformation frequently produced inflated false-positives and low sensitivity in simulations. In addition, Wilcoxon lacked the flexibility to accommodate complex study designs involving continuous traits or confounders. ANCOMBC2 also exhibited highly inflated FDRs in both permutations and simulations, with more false-positive discoveries in less abundant cell types.

Overall, LOCOM showed the best performance, followed by ANCOM-BC and LR+CLR, which together emerged as the top-performing methods among those evaluated (**Table 4**). LOCOM performed consistently well across most permutation and simulation settings and was robust to differential library sizes and correlations among cell groups. It also accommodates confounders and continuous traits, making it the most reliable and flexible option. However, LOCOM assumes that at least half of the cell groups are not differentially abundant, a common assumption among microbiome tools, and should therefore be interpreted cautiously when widespread compositional shifts are expected. ANCOM-BC also showed stable and competitive performance across most settings, although it exhibited persistent type I error inflation in the ILD datasets. LR+CLR is straightforward to implement and generally achieved high calibrated sensitivity, but its FDR control deteriorated under strong differential signals and its results may be sensitive to pseudocount selection. These complementary strengths and limitations suggest that no single method is uniformly optimal. Specifically, for binary-trait analyses, with or without confounders, we recommend considering LOCOM, ANCOM-BC, and LR+CLR. For continuous-trait analyses, which are not supported by many methods, LOCOM is a strong option. In general, we recommend applying more than one top-performing method and prioritizing findings that are consistently supported across methods, while remaining mindful of the limitations of each approach.

Prior to selecting a method, it is essential to assess the suitability of the dataset for differential composition analysis and to carefully preprocess the data to ensure reliable results. Firstly, based on our simulations with small sample sizes, we do not recommend conducting differential composition analysis on datasets with a total sample size of 10 or fewer, as all methods exhibited very low sensitivity in such settings. Secondly, data preprocessing play a crucial role in the differential composition analysis. While the standard of data filtering during preprocessing remains an open question, we empirically suggest retaining samples with at least 800 - 1000 cells and cell groups with a minimum prevalence of 10% - 20%, in line with established guidelines from differential abundance analysis in microbiome studies [14]. Empirically, we observed that samples with very low and sparse cell counts often lead to failure of some methods, such as convergence issues. Finally, since the differential composition methods evaluated in this study heavily depend on upstream cell type clustering and annotation, it is critical that researchers validate their clustering results and cell type annotations before proceeding to investigate compositional differences.

Several important considerations remain in our study. First, we encourage users to interpret the overall rankings in conjunction with detailed performance metrics for each method under specific settings. Our ranking system penalizes FDR inflation but not deflation, and only summarizes the relative performance of methods within each setting. Second, our evaluation primarily focused on binary and continuous traits, and the method performance under more complex designs, such as multi-group comparisons or repeated measures, remains to be investigated in future work. Third, all the methods benchmarked in this study are clustering-based, relying on counts of predefined cell types. In contrast, an emerging class of clustering-free differential abundance methods defines cell neighborhoods using gene expression data to detect local enrichment for specific conditions, without relying on discrete clustering [38, 39]. These methods may be more robust to clustering inaccuracy and can capture local subtle changes, though their performance can depend on neighborhood construction and may be less straightforward to interpret. We note that clustering-based and clustering-free approaches are fundamentally distinct. Clustering-based methods are well-suited for detecting changes in the abundance of well-defined cell types, whereas clustering-free methods are better equipped to reveal new or previously uncharacterized cellular groups associated with the condition. Given these distinct goals, we did not include clustering-free approaches in our benchmarking study, which have been evaluated elsewhere [39].

Despite the availability of differential composition methods in single-cell data, there remain multiple avenues for methodological refinement, particularly to address the unique characteristics of scRNA-seq. As noted earlier, clustering-based methods are inherently sensitive to clustering parameters and cell type annotation. One promising direction is to incorporate the uncertainty from cell type assignments into differential composition analysis. For example, DCATS offers the ability to account for potential misclassification by incorporating a similarity matrix among cell types [15]. However, since this feature requires an additional layer of input (gene expression) not used by other methods, it was not evaluated in our benchmarking. Similarly, scDC, which was not included due to its runtime inefficiency, captures cell type proportion uncertainty via bootstrap-based confidence intervals [40]. Another critical challenge is the potential bias in the single-cell experimental workflow [41, 42]. Steps in the wet-lab process can distort true cell type proportions. However, due to the lack of ground truth data for cell type composition, such biases are difficult to estimate and remain a persistent issue. Finally, as the focus of single-cell research shifts from broad cell types to finer subtypes and dynamic cell states, it becomes increasingly relevant to consider hierarchical or grouped structures in the analysis. Traditional approaches that apply uniform multiple testing correction across all subtypes may suffer from reduced power. In contrast, hierarchical or grouped testing strategies could enhance power by leveraging the natural grouping of subtypes under a common parent cell type.

## 4 Methods

### 4.1 Real Data Preprocessing

Cell group annotations at both the CT and CS levels for the AD, ILD, and OneK1K scRNA-seq datasets were obtained from the original publications. For each level of annotation, a sample-by-cell-group count matrix was first generated by counting the number of cells per cell group. Samples containing fewer than 800 cells were excluded from further analysis. Additionally, cell groups were filtered out if they had a proportion of zero greater than 0.8.

Binary disease status in the AD and ILD datasets and sex in the OneK1K dataset were used as the traits of interest. For the AD datasets, the binary disease status was derived from the final consensus cognitive diagnosis [43]. Specifically, values of 4 (AD with no other cause) and 5 (AD with an additional cause of cognitive impairment) were coded as TRUE, while all other values were coded as FALSE.

### 4.2 Method Implementation

#### LR + CLR and Wilcoxon + CLR

A pseudo-count of 0.5 was added to all cell counts to address zeros in the cell count data [44]. The counts were then converted to relative abundances, followed by CLR transformation. For each cell group, LR and Wilcoxon rank-sum tests were performed using the aov() and wilcox.test() functions in R, respectively, except in confounder or continuous-trait settings. In these settings, linear models were fitted using the lm() function and evaluated by F tests using the anova() function. In confounder settings, full models including the trait of interest and confounder terms were compared against reduced models including only the confounder. Resulting P-values were adjusted across cell groups using the BH procedure.

#### Quasi-binomial F test

Cell counts were converted to relative abundances and modeled for each cell group using a quasi-binomial generalized linear model with a logit link via the glm() function in R. F tests were performed using the anova() function. In confounder settings, full models including the trait of interest and confounder terms were compared against reduced models including only the confounder. Resulting P-values were adjusted across cell groups using the BH procedure.

#### DCATS

DCATS was evaluated with the R/Bioconductor package DCATS (v1.2.0, https://www.bioconductor.org/packages/release/bioc/html/DCATS.html) in R. For binary traits, we followed the package vignette to use dcats GLM() and set the parameter similarity mat to NULL. For continuous traits, we used the continuous variable in the design matrix and set base model in dcats GLM() to NULL.

#### scCODA

scCODA was evaluated with the Python package scCODA (v0.1.9, https://github.com/theislab/scCODA) following the package documentation.

#### sccomp

The sccomp R/Bioconductor package (v1.8.0) was applied following the vignette provided on Bioconductor: https://www.bioconductor.org/packages/release/bioc/html/sccomp.html. The parameter bimodal mean variability association was set to TRUE, as recommended in the documentation. Based on runtime messages, we set variational inference to FALSE to enable full Bayesian inference instead of variational Bayes. Since sccomp includes an optional outlier removal step, we implemented the method in two ways: (i) *sccomp* (without outlier removal) and (ii) *sccomp-or* (with outlier removal).

#### propeller

Propeller, included in the speckle R/Bioconductor (v1.4.0, https://www. bioconductor.org/packages/release/bioc/html/speckle.html) package, was implemented following the original publication [16]. The propeller.anova() function was used when confounder adjustment was required, while the propeller() function was used for all other analyses involving binary traits.

#### LOCOM

The LOCOM R package (v2.0) was applied using the locom() function, following the instructions provided in the LOCOM GitHub repository https://github.com/yijuanhu/ LOCOM.

#### ANCOM-BC and ANCOM-BC2

Both methods were applied using the ANCOMBC R/Bioconductor package (v2.6.0, https://www.bioconductor.org/packages/release/bioc/html/ANCOMBC.html). The count matrix and design matrix were converted into TreeSummarizedExperiment objects and used as input for the ancombc() and ancombc2() functions. Parameters lib cut and prv cut were both set to 0, as the input data had been pre-processed. P-values were adjusted using the BH procedure, and a nominal FDR threshold of 0.1 was specified via the alpha parameter.

#### DESeq2

DESeq2 R/Bioconductor package (v1.44.0, https://bioconductor.org/packages/release/bioc/html/DESeq2.html) was used. Cell count matrices were converted to DESeqDataSet objects, and differential composition was assessed using the DESeq() function. The parameter sfType was set to poscount.

### 4.3 Simulator Evaluation

Four compositional data simulators, including MIDASim (parametric mode), sparseDOSSA, the simulation function of sccomp, and the DM model were applied to the control groups of AD CS, AD CT, ILD CS and ILD CT, as well as to female samples from OneK1K CS, generating 100 replicates per template and per simulator. To evaluate the global similarity between real and simulated samples, we applied principal coordinates analysis (PCoA; R package phyloseq [45]) and uniform manifold approximation and projection (UMAP; R package umap [46]) on Bray-Curtis distances. Permutational multivariate analysis of variance (PERMANOVA) was used to test for differences between simulated and observed samples via the R package vegan [47, 48] across 100 replicates. At the cell-group level, we assessed key cell-group features, including the proportion of zero, mean relative abundance, log_10_ total counts per cell group, and overdispersion (variance-to-mean ratio), and compared them between simulated and observed samples.

### 4.4 Simulation Details

We used the control group from the AD and ILD datasets and the female group from the OneK1K dataset as templates for simulation. To generate data under the global null hypothesis, we fitted a simulation model using the template samples and used this model to simulate data for both trait groups.

To generate data under the alternative hypothesis, we randomly selected 20% of cell groups as causal groups in each simulation run. For each causal group, an effect magnitude was sampled from a Beta(2, 2) distribution, and its direction was assigned to be positive with probability 0.8 and negative with probability 0.2. The baseline mean relative abundance of each causal group was modified according to the exponential of its effect size, followed by renormalization across all cell groups to preserve the compositional constraint. We considered multiple signal strengths by scaling the simulated effect sizes by 0.25, 0.50, 0.75, and 1 for the AD and OneK1K scenarios, and by 0.75, 1.5, 2.25, and 3 for the ILD scenarios. Samples in the control group were simulated from the model fitted to the template samples, while samples in the case group were simulated from an adjusted model incorporating the true effect sizes. Model fitting and adjustment were performed using MIDASim.setup() and MIDASim.modify() from the R package MIDASim, and the corresponding functions in sparseDOSSA. The simulation module included seven settings, summarized below:

#### The default setting

This setting simulates data with observed library sizes and observed cell-group correlations. It does not incorporate confounders, considers a binary trait, employs the MIDASim simulator, and uses the sample sizes of the real datasets.

#### Non-differential library sizes

Compared to the default setting, this setting removes the difference in library sizes as follows: Library sizes from the control (or female) group were used for both sample groups to remove differences in library sizes. If the case/male group had more samples than the control/female group, library sizes were sampled with replacement from the control/female group.

#### No correlations

This setting further removed observed correlations between cell groups compared to the setting with non-differential library sizes. We used a custom function MIDASim no corr() (adapted from the MIDASim function MIDASim()) to generate cell count data without cell-group correlations.

#### Confounders

This setting added a binary confounder *C_i_* to the default setting. Specifically, *C_i_* followed a Bernoulli distribution with a probability of 0.2 for samples with the binary trait of interest *T_i_* = 0 and 0.8 for samples with *T_i_* = 1. Confounding signals were added to the top 30% of the most abundant cell groups with differential signals and the top 20% without differential signals. The effect sizes of the confounder in selected cell groups were set to that of the binary trait plus one, and the mean relative abundances were subsequently normalized to ensure that they summed to 1 across all cell groups.

#### Continuous traits

A continuous trait was drawn from a standard normal distribution and scaled to have the same norm as the binary trait. The other simulation parameters in this setting are the same as the default setting.

#### An alternative simulator

As sparseDOSSA controls library sizes by specifying the population’s median library sizes and thus does not support specifying differential library sizes across groups, we use it to reproduce the results in the default setting with non-differential library sizes and observed correlations.

#### Small sample sizes

This setting simulates data with sample sizes of 10, 30, 50, and 70, using the templates from the five scenarios. An equal number of samples is assigned to each group of the binary trait, and library sizes are randomly sub-sampled by group from the real datasets. For each combination of sample size and scenario, it simulates null datasets with effect sizes equal to 0 and alternative datasets with effect sizes estimated from real data.

### 4.5 Benchmarking Metrics

We estimated type I error as the proportion of replicates in which the nominal p-value was smaller than 0.05. We estimated the empirical FDR as the average proportion of false discoveries among all discoveries. Under the global null hypothesis, we further estimated a confidence interval for the empirical FDR at the nominal level of 0.1 using a binomial distribution, assuming

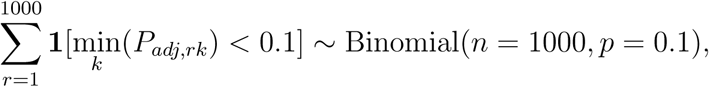

where *P_adj_* denotes the BH-adjusted p-value, *r* denotes the permutation or simulation replicate, and *k* denotes the cell group. This yielded an exact binomial confidence interval of [0.08, 0.12].

We estimated sensitivity as the average proportion of true discoveries among all true associations. As sensitivity tends to be higher for methods with inflated FDR, the direct comparison of sensitivity without accounting for FDR is unfair to methods with appropriate FDR control. To address this, we calibrate the sensitivity by reclaiming significance after adjusting the empirical FDR of all methods to a common target level of 0.1. Specifically, we first ranked all BH-adjusted p-values across 1,000 simulation replicates in ascending order. Starting from the smallest value, we sequentially considered each as a potential significance threshold and estimated empirical FDR at this threshold. We continued this process until identifying the largest value for which the empirical FDR was equal to 0.1, or as close as possible without exceeding it. This threshold was then used to claim significance, based on which we computed sensitivity, referred to as calibrated sensitivity. As sccomp is prohibitively slow and the calibration requires evaluation on 1,000 data replicates, we did not compute calibrated sensitivity for sccomp and sccomp-or.

## 5 Declarations

### 5.1 Ethics Approval, Consent to Participate, and Consent for Publication

Not applicable.

### 5.2 Data Availability

The snRNA-seq data from the ROSMAP study [1] are available on Synapse under accession code syn53366818. The scRNA-seq data for the ILD study [31] can be downloaded from the Gene Expression Omnibus (GEO) under accession number GSE227136. The OneK1K cohort cell-by-gene data [2] were downloaded from https://datasets.cellxgene.cziscience.com/e1239e77-732c-4dc6-9a4c-c9bb72607c61.rds.

### 5.3 Competing Interests

The authors declare no competing interests.

### 5.4 Funding

This work was supported in part by the National Institutes of Health award R01GM141074.

### 5.5 Author Contribution

W.W. implemented the methods, conducted the benchmarking analyses, and prepared all figures and supplementary materials. W.W. and M.H. developed the code. W.W., M.H., Y.H., and C.S. wrote the manuscript. Y.H. and C.S. conceived and supervised the study. All authors reviewed and approved the final manuscript.

### 5.6 Code Availability

All custom code necessary to reproduce the simulations, permutation analyses, real-data analyses, and figures and tables presented in this study is provided as a ZIP archive for editors and reviewers. The final version of the code will be archived in a permanent public repository upon publication.

## Notes

### Competing Interest Statement

The authors have declared no competing interest.

