## Supplementary Information for "Benchmarking Statistical Methods for Differential Composition Analysis in Population-Scale Single-Cell Data"

#### Contents


**1 Supplementary Figures** **2**

**2 Supplementary Tables** **31**

### 1 Supplementary Figures

a

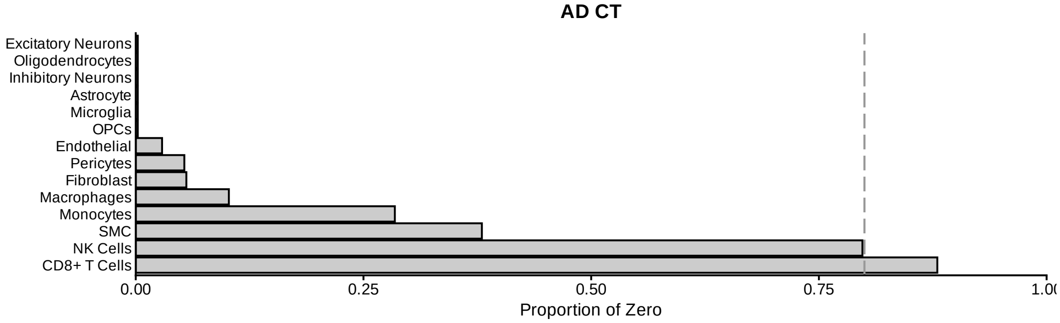


b

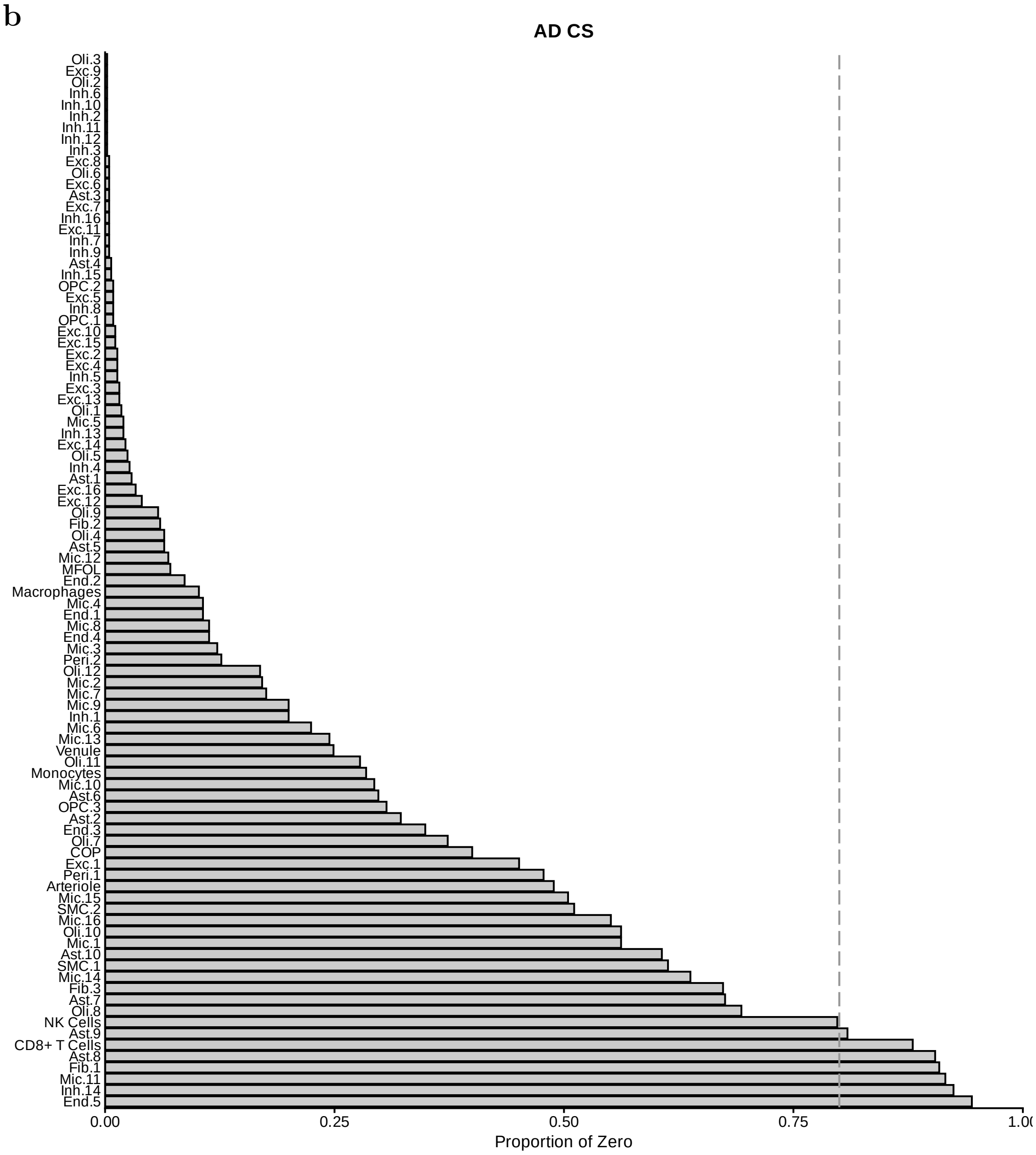

**c**

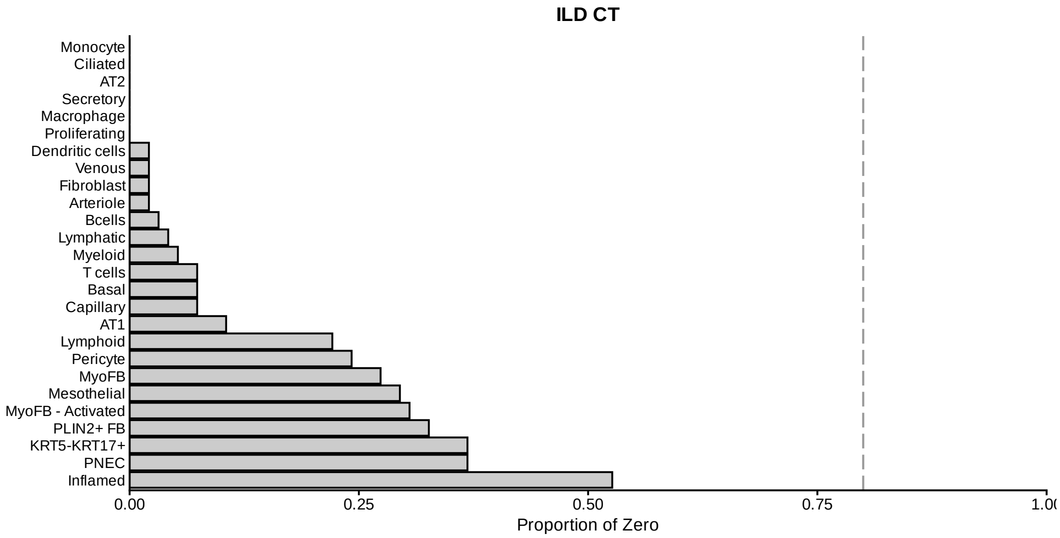
ILD CT
Monocyte
Ciliated
AT2
Secretory
Macrophage
Proliferating
Dendritic cells
Venous
Fibroblast
Arteriole
Bcells
Lymphatic
Myeloid
T cells
Basal
Capillary
AT1
Lymphoid
Pericyte
MyoFB
Mesothelial
MyoFB - Activated
PLIN2+ FB
KRT5-KRT17+
PNEC
Inflamed
0.00
0.25
0.50
0.75
Proportion of Zero

**d**

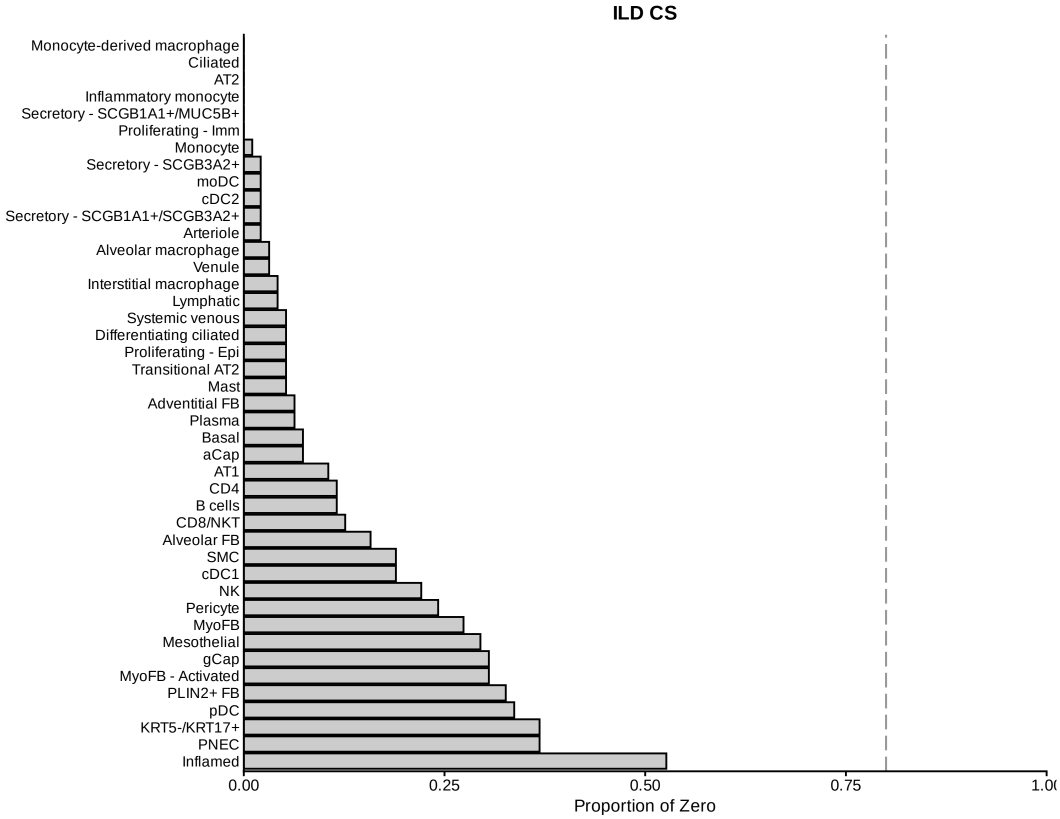
ILD CS
Monocyte-derived macrophage
Ciliated
AT2
Inflammatory monocyte
Secretory - SCGB1A1+/MUC5B+
Proliferating - Imm
Monocyte
Secretory - SCGB3A2+
moDC
cDC2
Secretory - SCGB1A1+/SCGB3A2+
Arteriole
Alveolar macrophage
Venule
Interstitial macrophage
Lymphatic
Systemic venous
Differentiating ciliated
Proliferating - Epi
Transitional AT2
Mast
Adventitial FB
Plasma
Basal
aCap
AT1
CD4
B cells
CD8/NKT
Alveolar FB
SMC
cDC1
NK
Pericyte
MyoFB
Mesothelial
gCap
MyoFB - Activated
PLIN2+ FB
pDC
KRT5-/KRT17+
PNEC
Inflamed
0.00
0.25
0.50
0.75
Proportion of Zero

**e**

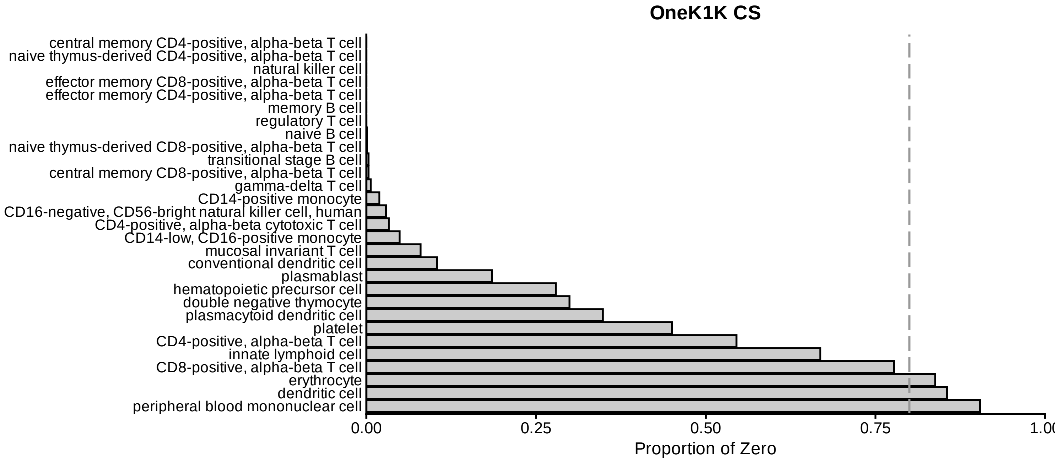


**Supplementary Fig. S1:** Proportion of zeros for each cell group in the five real datasets (**a**–**e**).

**a**

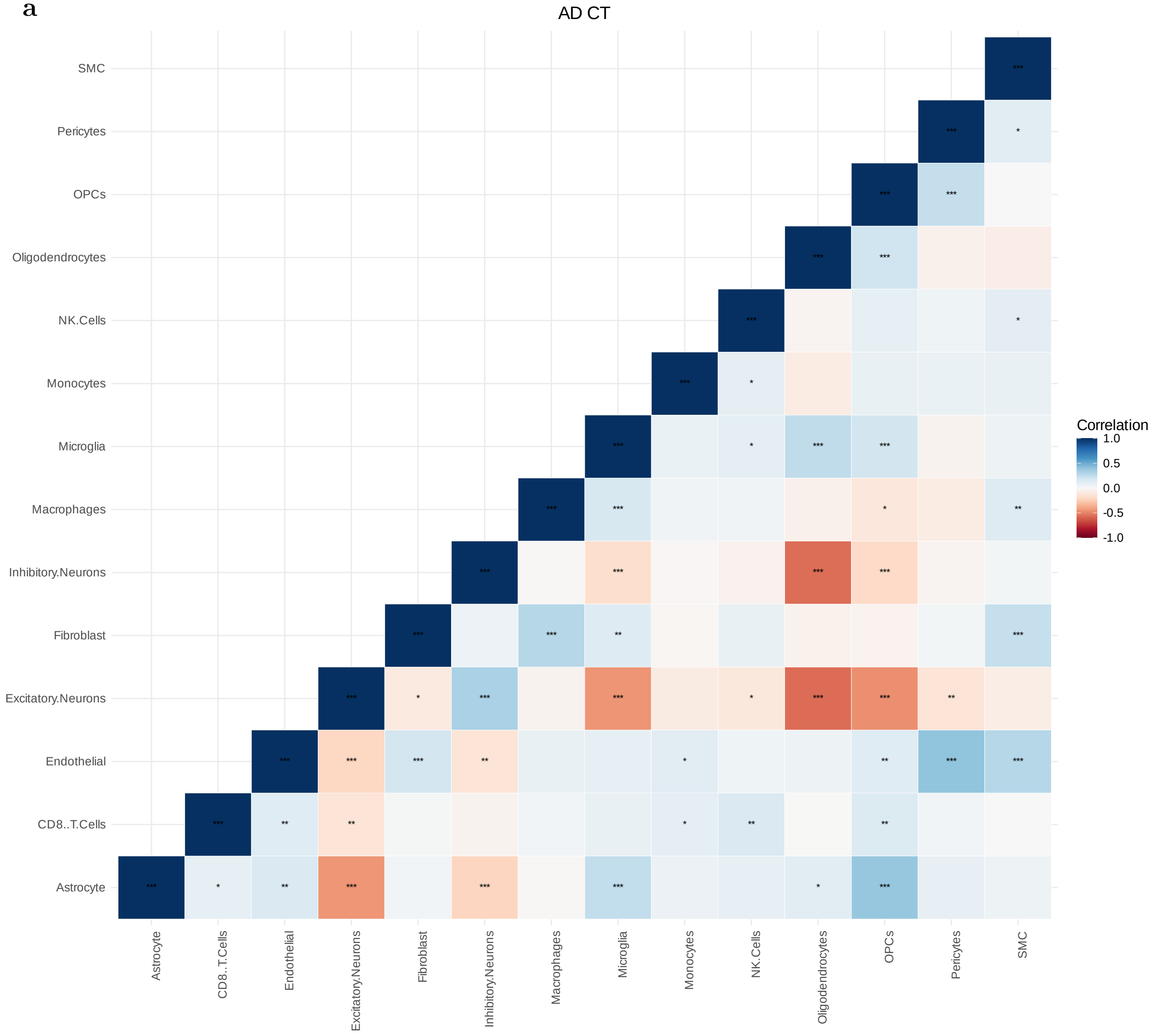
AD CT
SMC
Pericytes
OPCs
Oligodendrocytes
NK.Cells
Monocytes
Microglia
Macrophages
Inhibitory.Neurons
Fibroblast
Excitatory.Neurons
Endothelial
CD8..T.Cells
Astrocyte
Correlation
1.0
0.5
0.0
-0.5
-1.0

**b**

AD CS

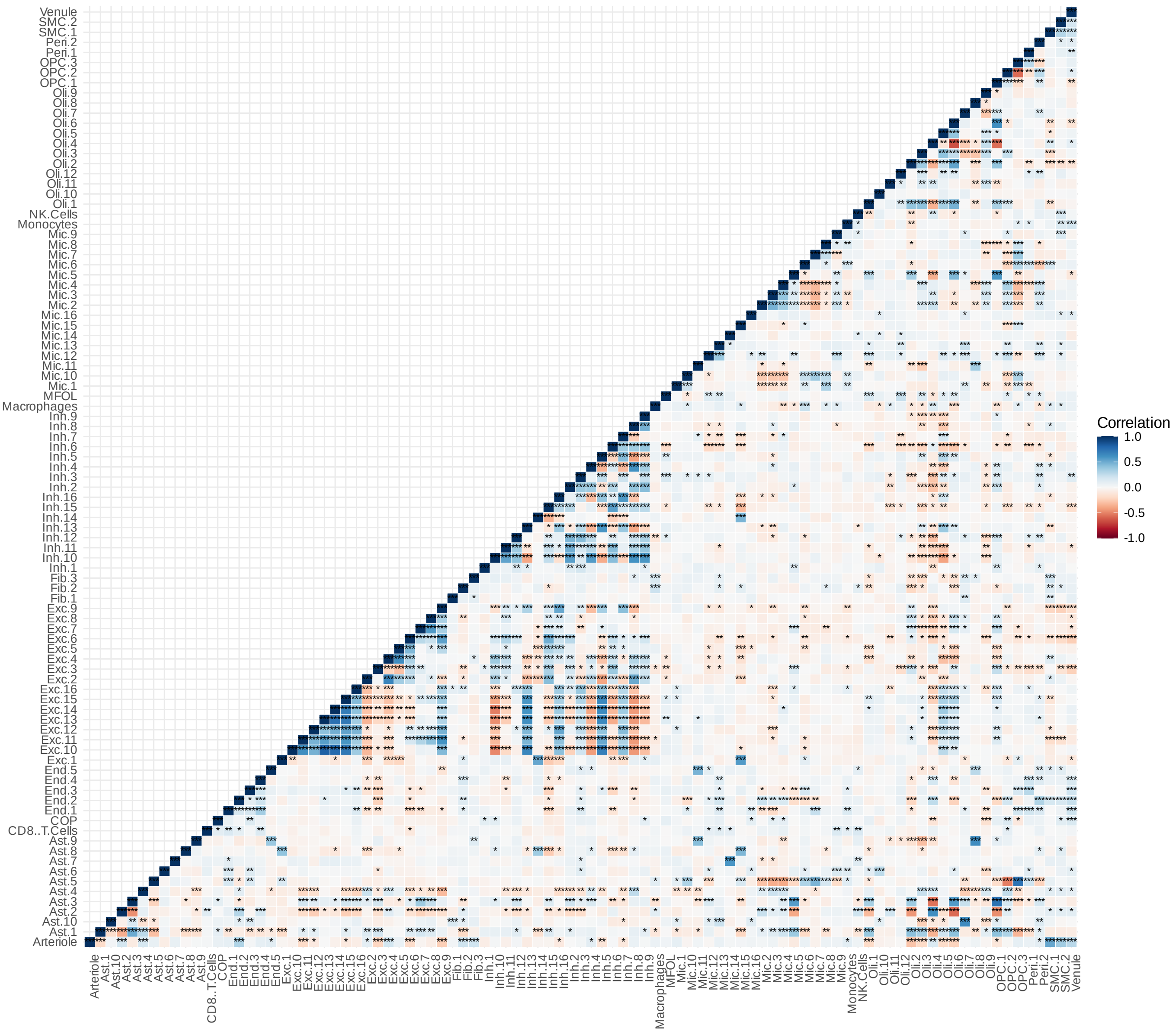
Correlation
1.0
0.5
0.0
-0.5
-1.0
Venule
SMC.2
SMC.1
Peri.2
Peri.1
OPC.3
OPC.2
OPC.1
Oli.9
Oli.8
Oli.7
Oli.6
Oli.5
Oli.4
Oli.3
Oli.2
Oli.12
Oli.11
Oli.10
Oli.1
NK.Cells
Monocytes
Mic.9
Mic.8
Mic.7
Mic.6
Mic.5
Mic.4
Mic.3
Mic.2
Mic.16
Mic.15
Mic.14
Mic.13
Mic.12
Mic.11
Mic.10
Mic.1
MFOL
Macrophages
Inh.9
Inh.8
Inh.7
Inh.6
Inh.5
Inh.4
Inh.3
Inh.2
Inh.16
Inh.15
Inh.14
Inh.13
Inh.12
Inh.11
Inh.10
Inh.1
Fib.3
Fib.2
Fib.1
Exc.9
Exc.8
Exc.7
Exc.6
Exc.5
Exc.4
Exc.3
Exc.2
Exc.16
Exc.15
Exc.14
Exc.13
Exc.12
Exc.11
Exc.10
Exc.1
End.5
End.4
End.3
End.2
End.1
COP
CD8..T.Cells
Ast.9
Ast.8
Ast.7
Ast.6
Ast.5
Ast.4
Ast.3
Ast.2
Ast.10
Ast.1
Arteriole

**c**

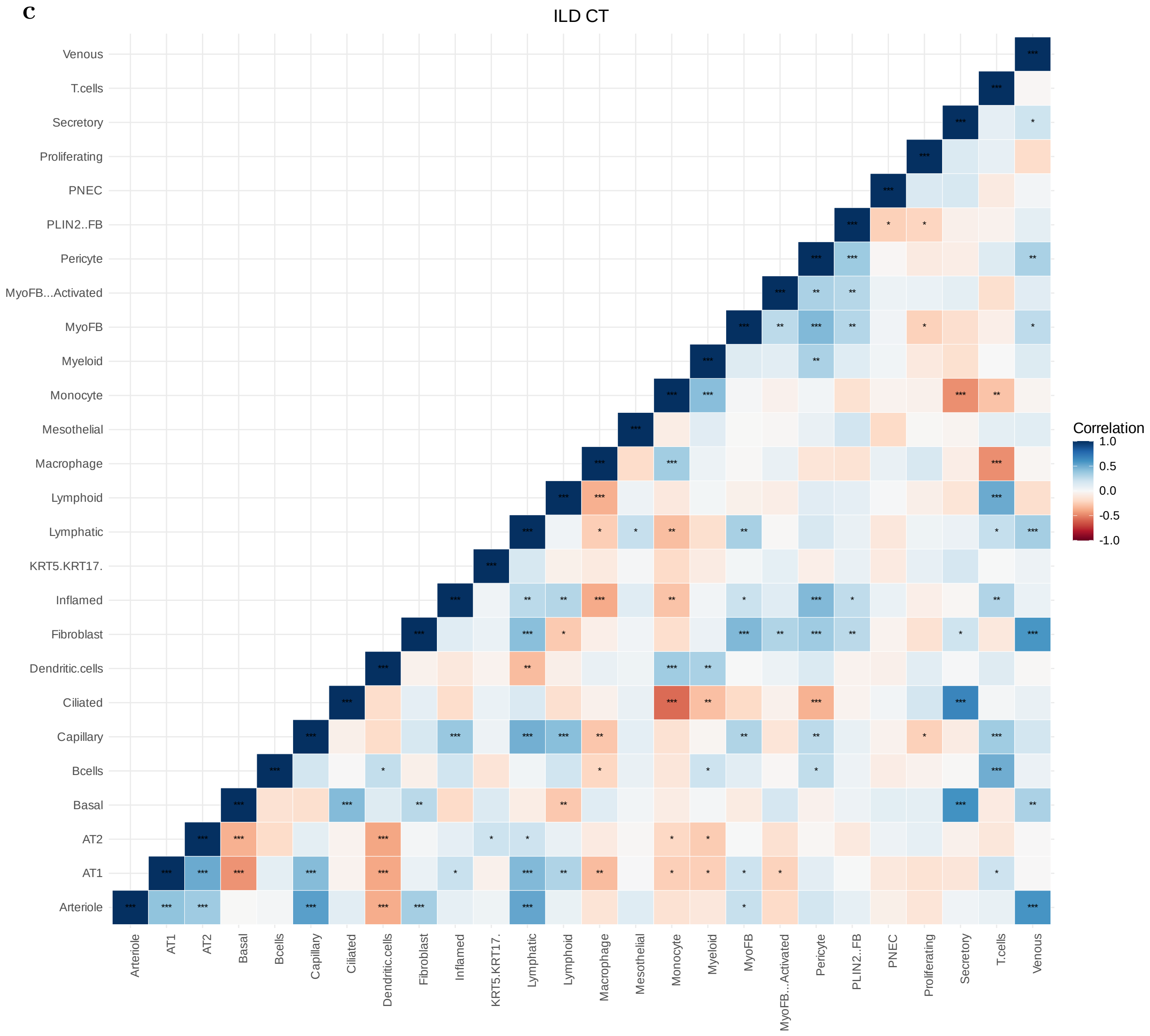
ILD CT
Venous
T.cells
Secretory
Proliferating
PNEC
PLIN2..FB
Pericyte
MyoFB...Activated
MyoFB
Myeloid
Monocyte
Mesothelial
Macrophage
Lymphoid
Lymphatic
KRT5.KRT17.
Inflamed
Fibroblast
Dendritic.cells
Ciliated
Capillary
Bcells
Basal
AT2
AT1
Arteriole
Correlation
1.0
0.5
0.0
-0.5
-1.0

**d**

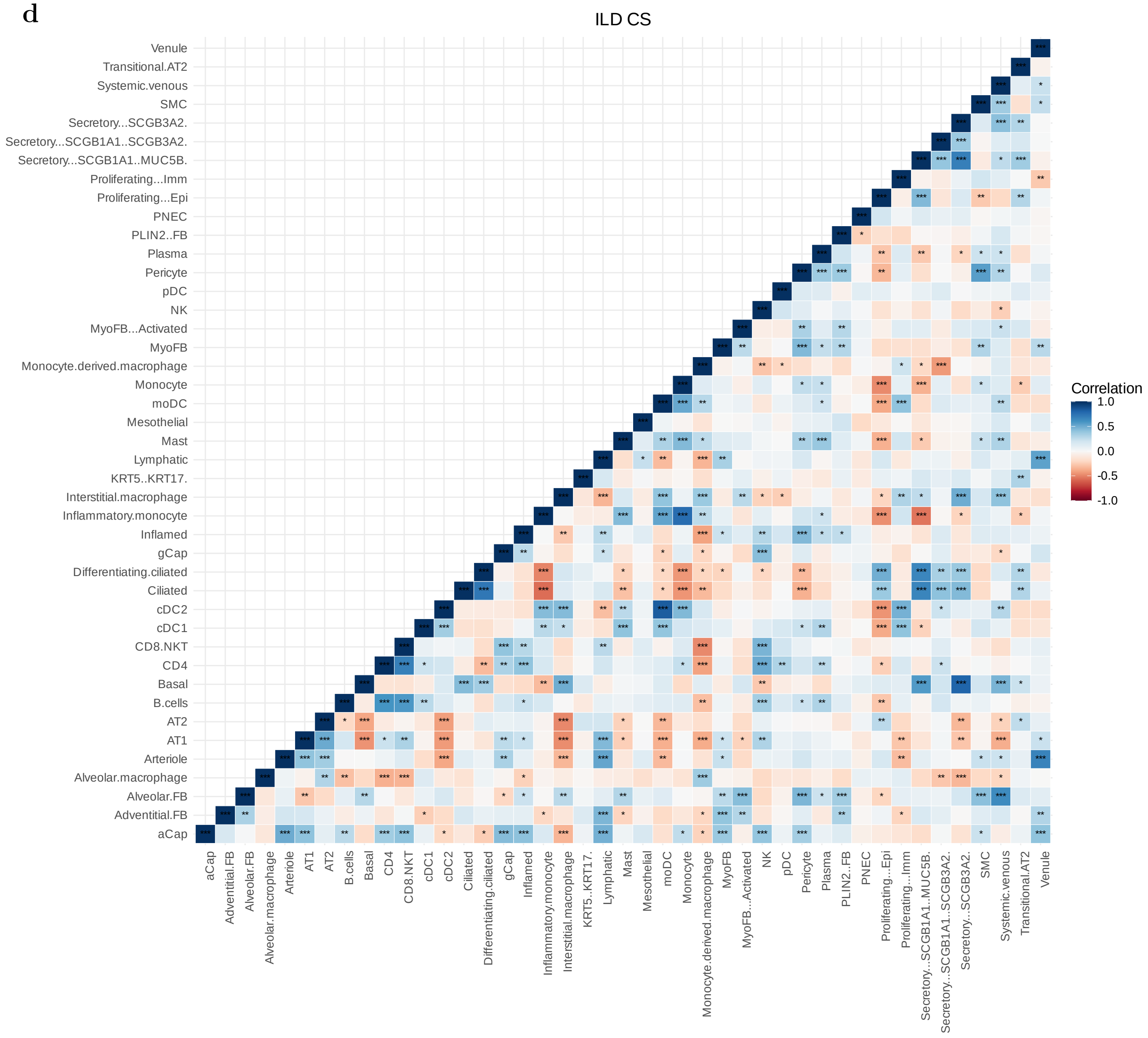
ILD CS
Venule
Transitional.AT2
Systemic.venous
SMC
Secretory...SCGB3A2.
Secretory...SCGB1A1..SCGB3A2.
Secretory...SCGB1A1..MUC5B.
Proliferating...Imm
Proliferating...Epi
PNEC
PLIN2..FB
Plasma
Pericyte
pDC
NK
MyoFB...Activated
MyoFB
Monocyte.derived.macrophage
Monocyte
moDC
Mesothelial
Mast
Lymphatic
KRT5..KRT17.
Interstitial.macrophage
Inflammatory.monocyte
Inflamed
gCap
Differentiating.ciliated
Ciliated
cDC2
cDC1
CD8.NKT
CD4
Basal
B.cells
AT2
AT1
Arteriole
Alveolar.macrophage
Alveolar.FB
Adventitial.FB
aCap
Correlation
1.0
0.5
0.0
-0.5
-1.0

**e**

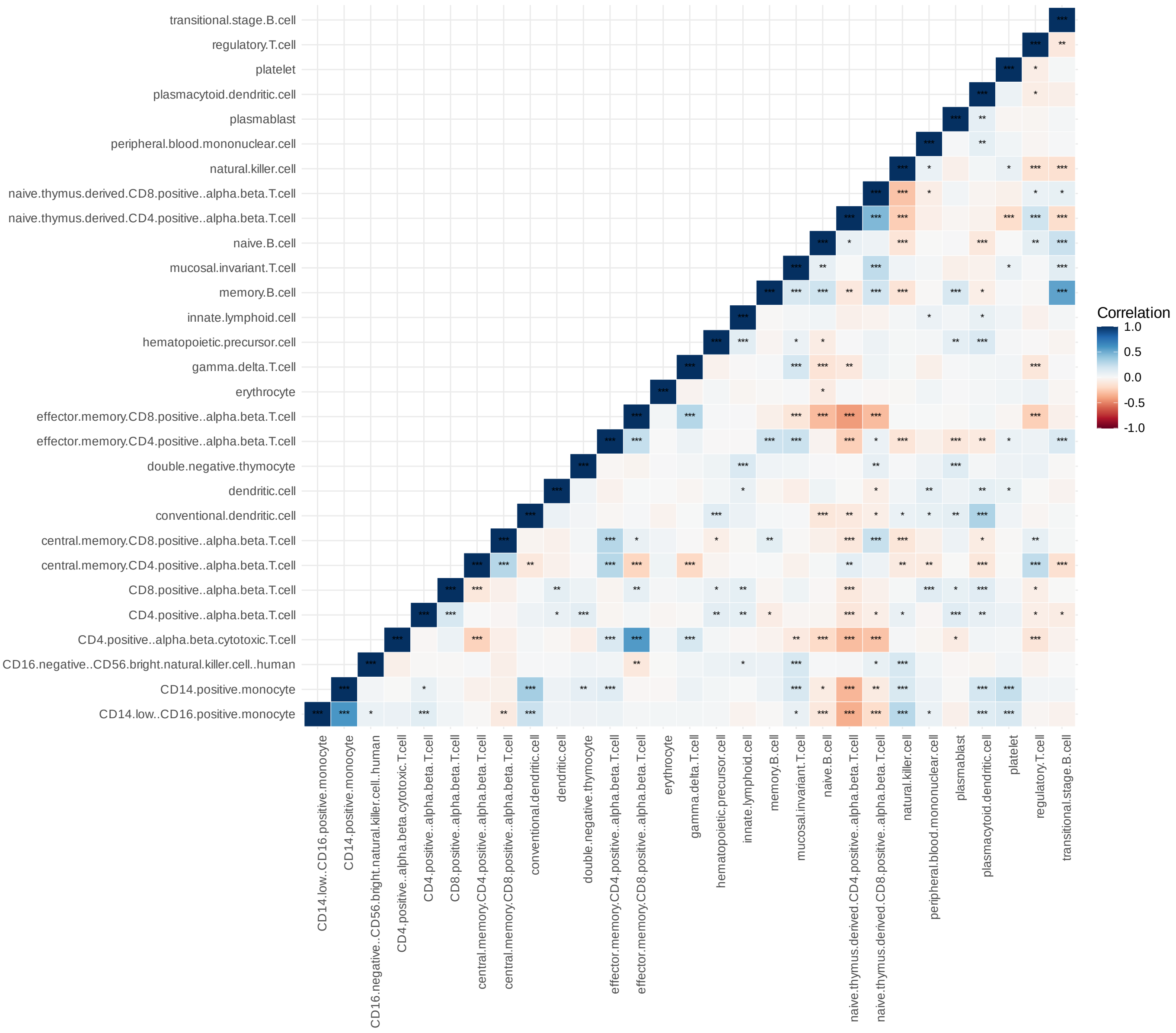


**Supplementary Fig. S2:** Correlations between cell groups in the five real datasets (**a**–**e**). Significance is indicated as *:P $<$ 0.05; **:P$<$ 0.01; and ***:P$<$ 0.001.

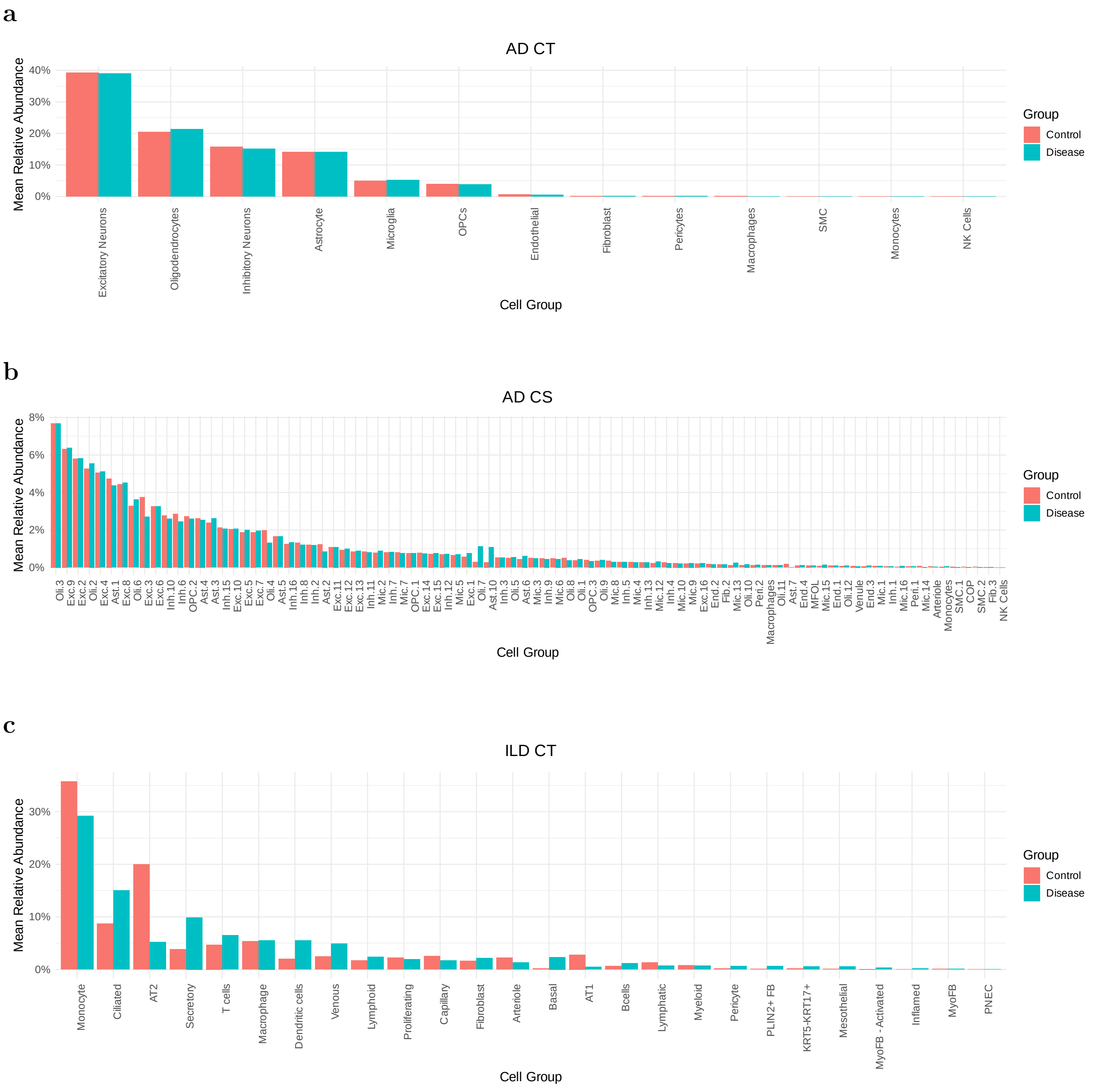
a
AD CT
Mean Relative Abundance
40%
30%
20%
10%
0%
Excitatory Neurons
Oligodendrocytes
Inhibitory Neurons
Astrocyte
Microglia
OPCs
Endothelial
Fibroblast
Pericytes
Macrophages
SMC
Monocytes
NK Cells
Cell Group
Group
Control
Disease
b
AD CS
Mean Relative Abundance
8%
6%
4%
2%
0%
Oli.3 Exc.9 Exc.2 Oli.2 Exc.4 Ast.1 Exc.8 Oli.6 Exc.3 Exc.6 Inh.10 Inh.6 OPC.2 Ast.4 Ast.3 Inh.15 Exc.10 Exc.5 Exc.7 Oli.4 Ast.5 Inh.16 Inh.8 Inh.2 Ast.2 Exc.11 Exc.12 Exc.13 Inh.11 Mic.2 Inh.7 Mic.7 OPC.1 Exc.14 Exc.15 Inh.12 Mic.5 Exc.1 Oli.7 Ast.10 Inh.3 Oli.5 Mic.3 Ast.6 Inh.9 Mic.6 Oli.8 Oli.1 OPC.3 Oli.9 Mic.8 Inh.5 Mic.4 Inh.13 Mic.12 Inh.4 Mic.10 Mic.9 Exc.16 End.2 Fib.2 Mic.13 Oli.10 Peri.2 Macrophages Oli.11 Ast.7 End.4 MFOL Mic.15 End.1 Oli.12 Venule End.3 Mic.1 Inh.1 Mic.16 Peri.1 Mic.14 Arteriole Monocytes SMC.1 COP SMC.2 Fib.3 NK Cells
Cell Group
Group
Control
Disease
c
ILD CT
Mean Relative Abundance
30%
20%
10%
0%
Monocyte
Ciliated
AT2
Secretory
T cells
Macrophage
Dendritic cells
Venous
Lymphoid
Proliferating
Capillary
Fibroblast
Arteriole
Basal
AT1
Bcells
Lymphatic
Myeloid
Pericyte
PLIN2+ FB
KRT5-KRT17+
Mesothelial
MyoFB - Activated
Inflamed
MyoFB
PNEC
Cell Group
Group
Control
Disease

**d**

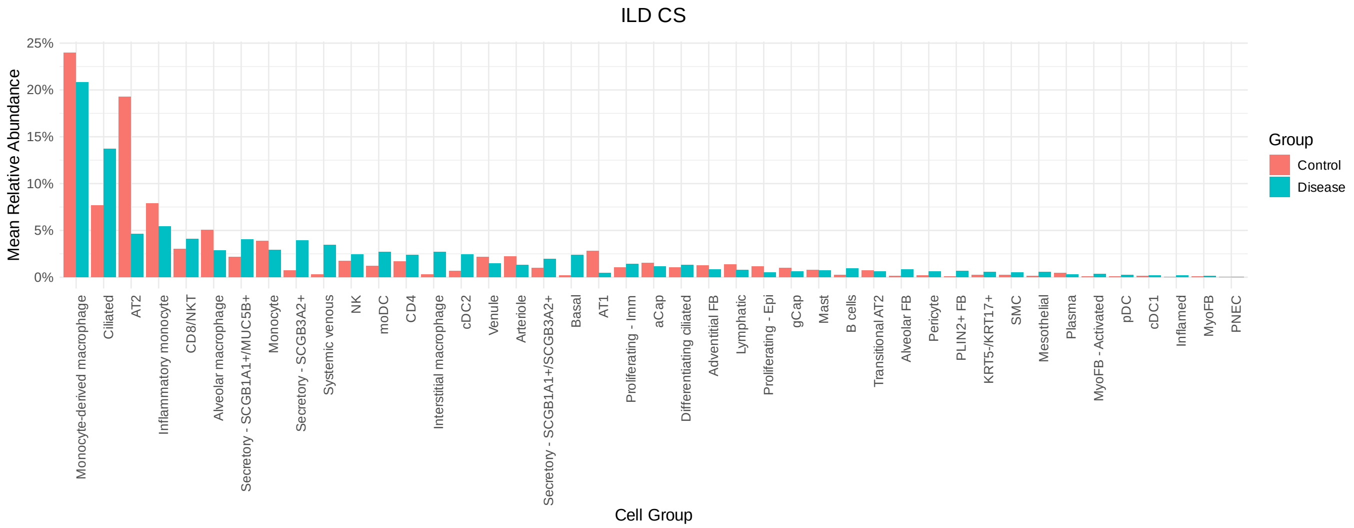


**e**

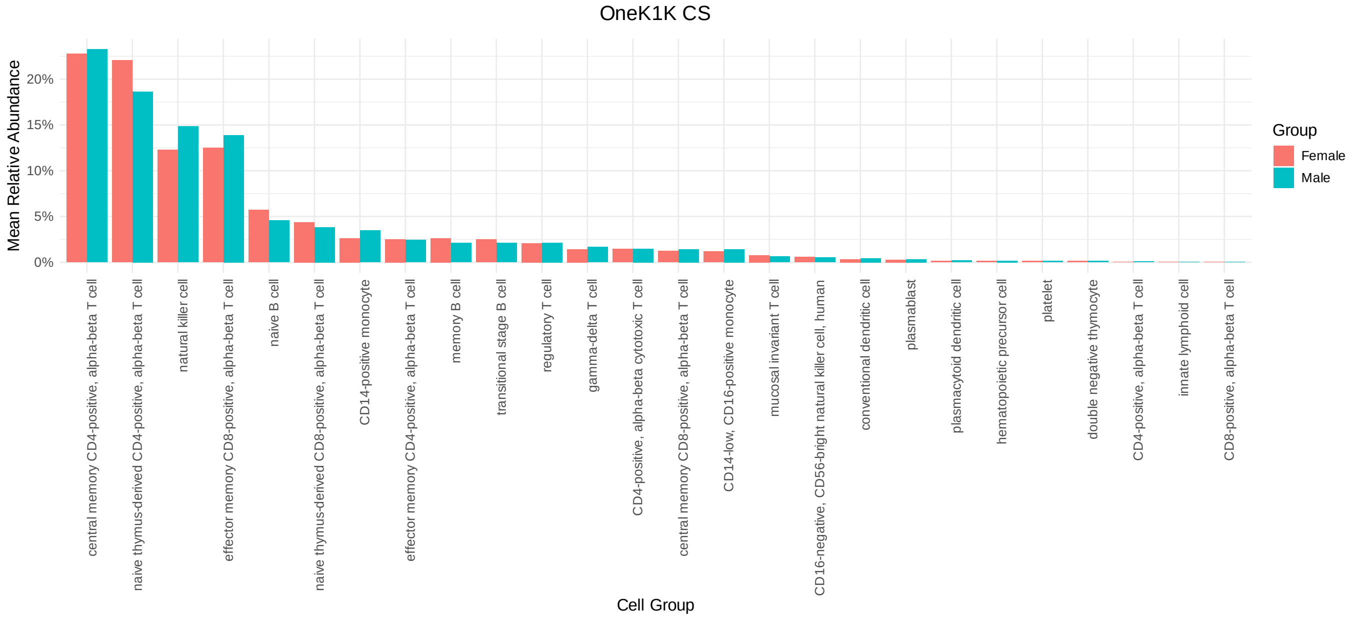


**Supplementary Fig. S3:** Mean relative abundances of cell groups by condition in the five real datasets (**a-e**).

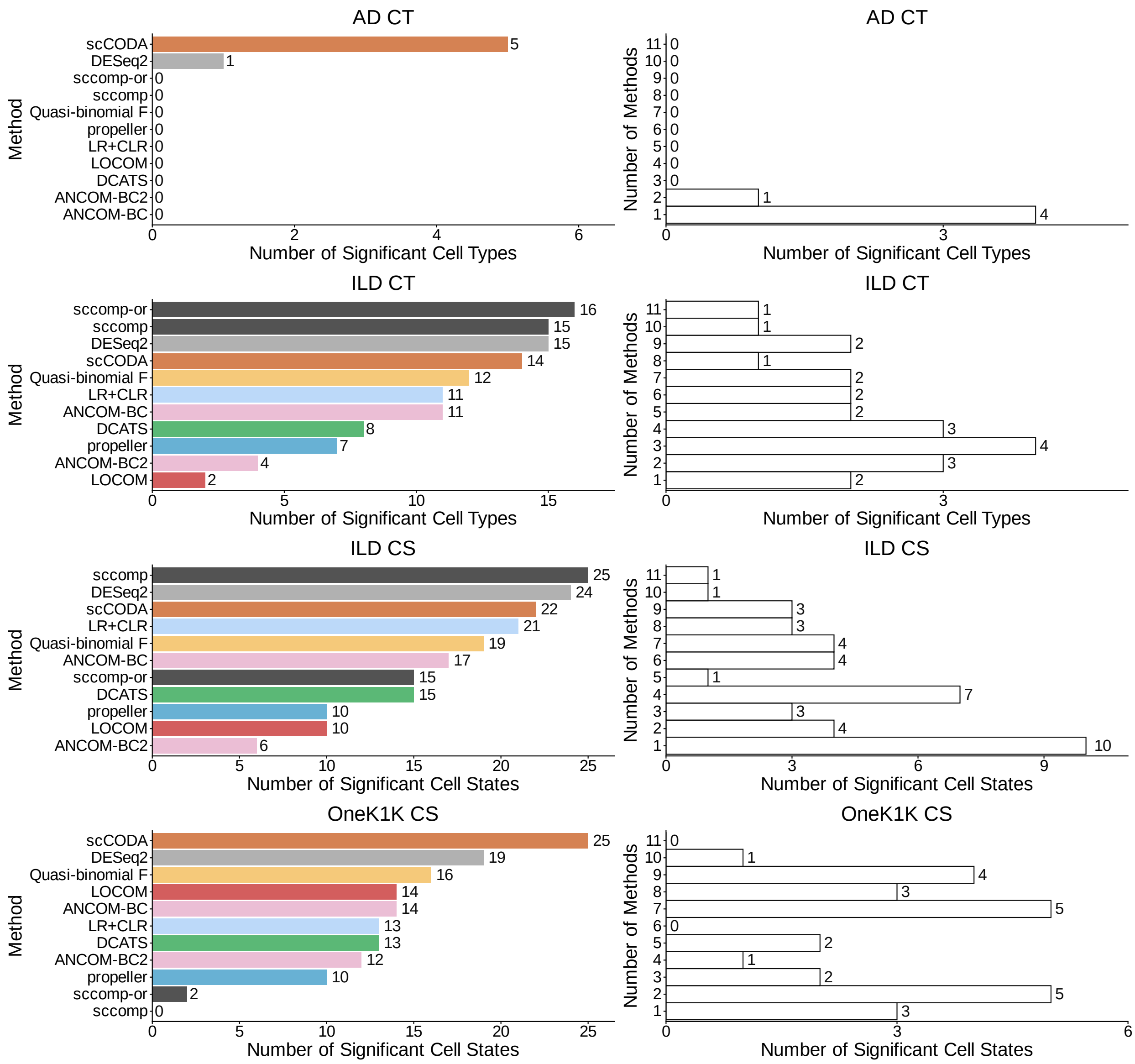


**Supplementary Fig. S4:** Analysis results of four real datasets with covariate adjustment. Left panel: number of cell states detected by different methods. Right panel: number of "shared" detected cell states across varying numbers of methods.

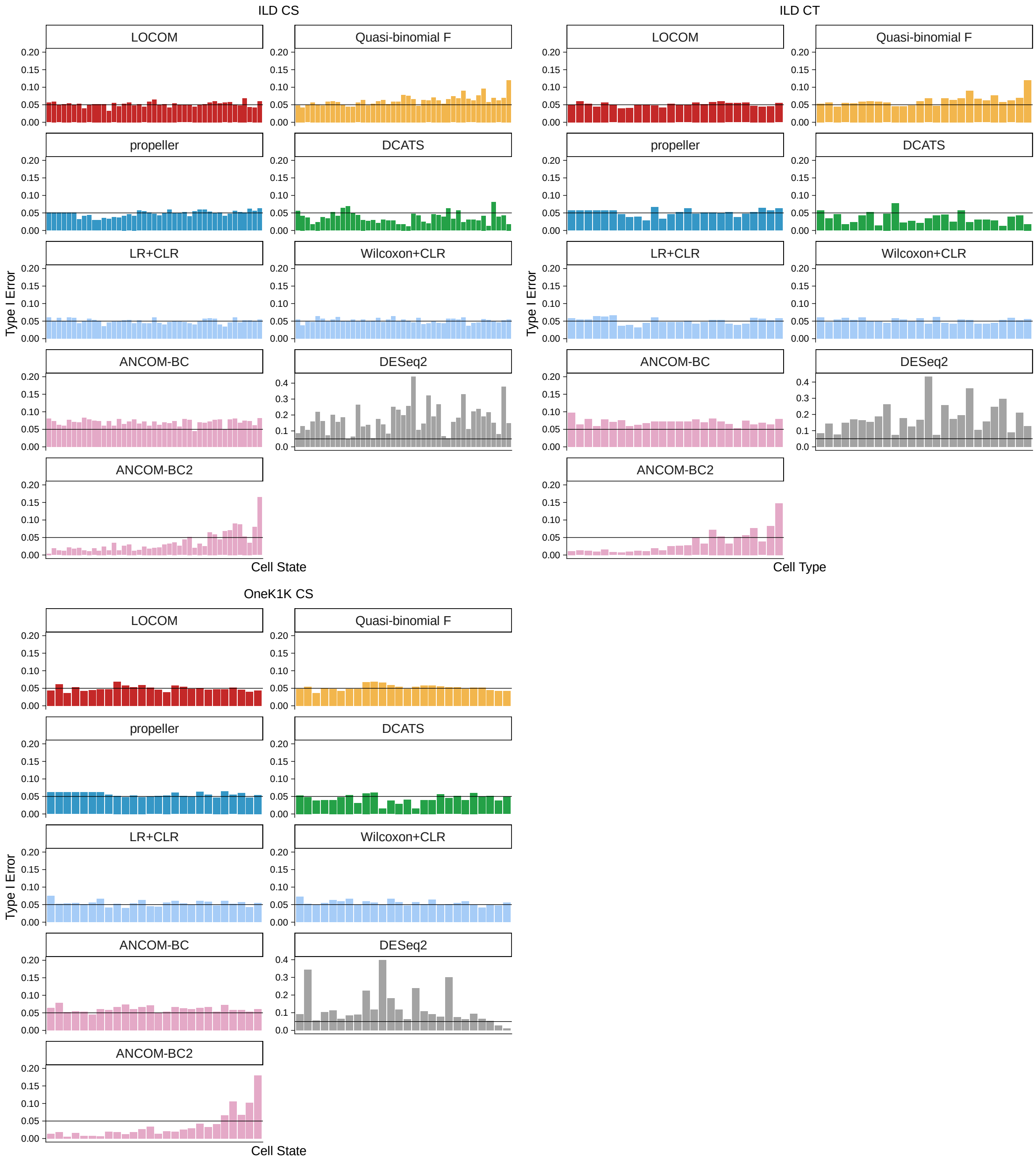


**Supplementary Fig. S5:** Type I error rates by cell group in analysis of permuted replicates of the ILD CS, ILD CT, and OneK1K datasets. The x-axes denote cell types and cell states in ascending order of the proportion of zero.

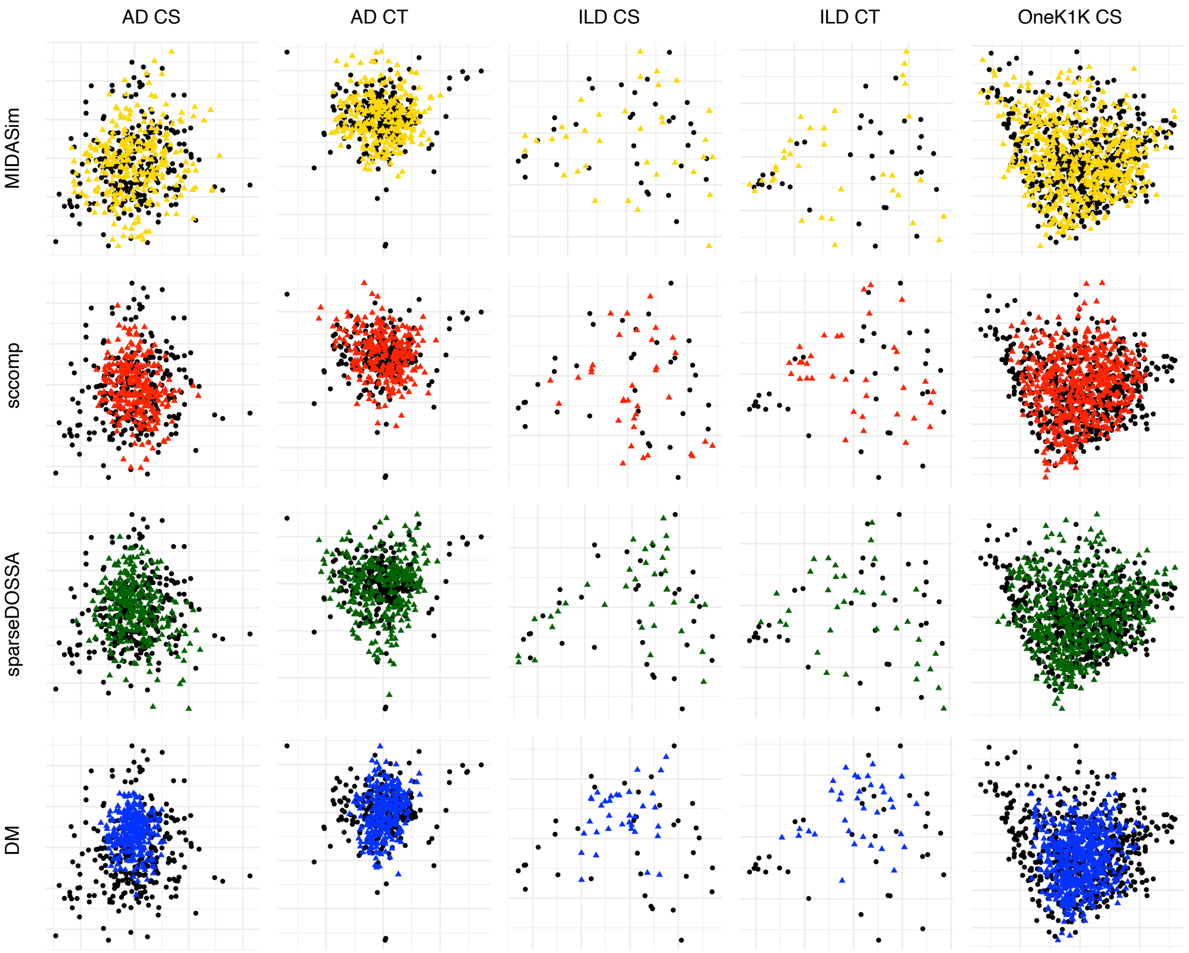


**Supplementary Fig. S6:** Principal coordinates plots (PCoA) of observed and simulated samples by four simulators across five template datasets. Black points denote observed samples; colored points denote simulated samples.

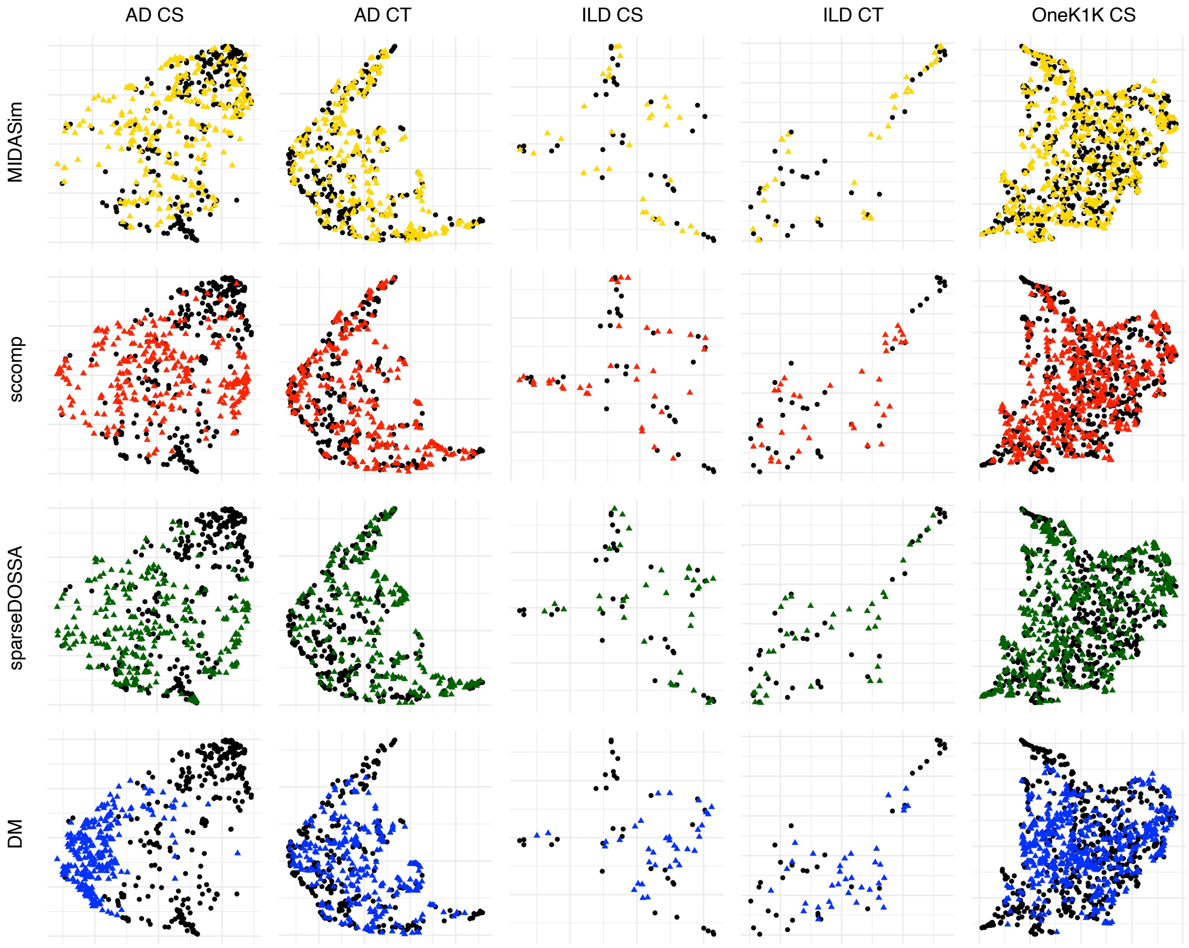


**Supplementary Fig. S7:** Uniform Manifold Approximation and Projection (UMAP) plots of observed and simulated samples by four simulators across five template datasets. Black points denote observed samples; colored points denote simulated samples.

**a**

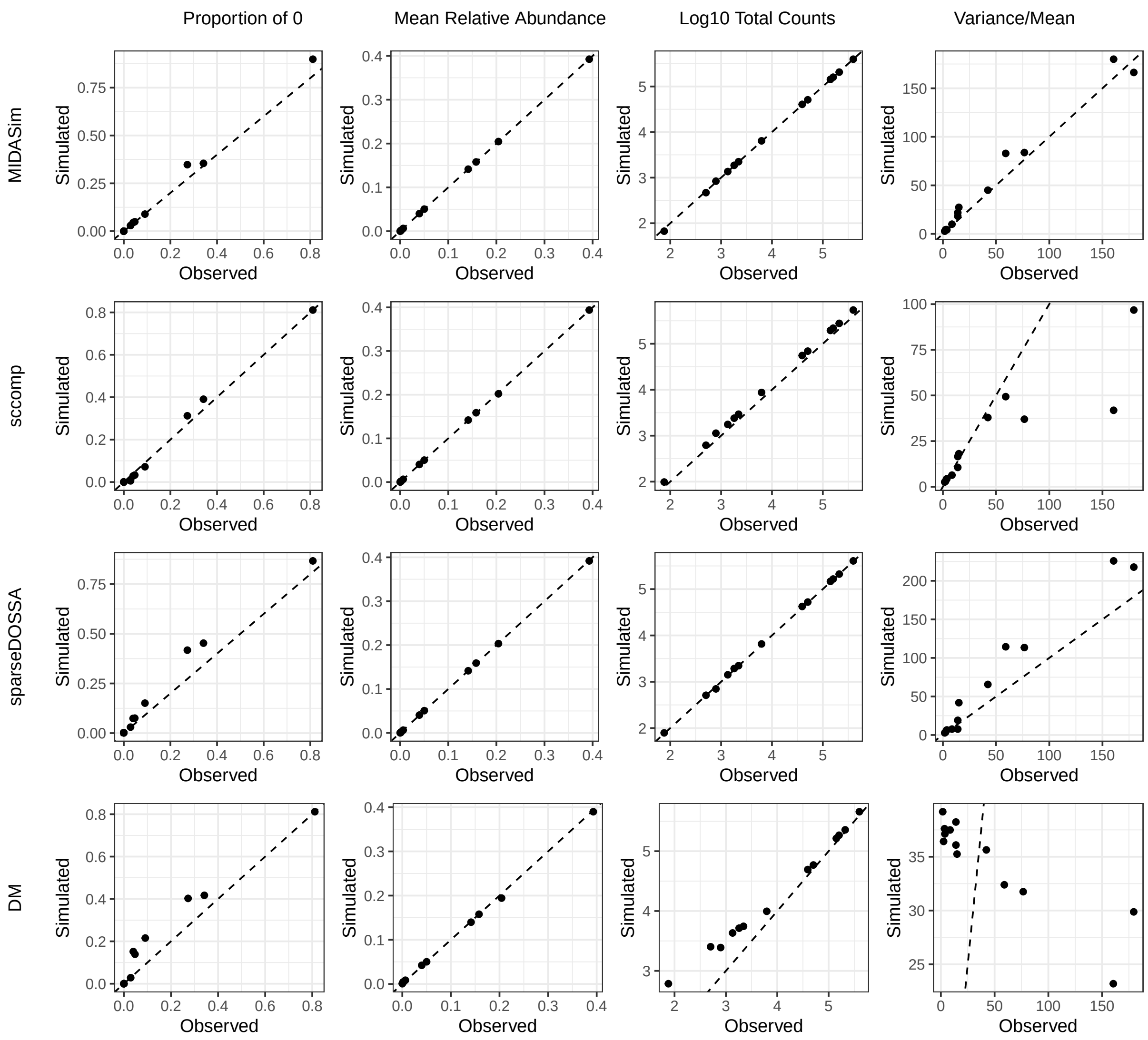

Proportion of 0
Mean Relative Abundance
Log10 Total Counts
Variance/Mean
MIDASim
sccomp
sparseDOSSA
DM
Simulated
Observed

**b**

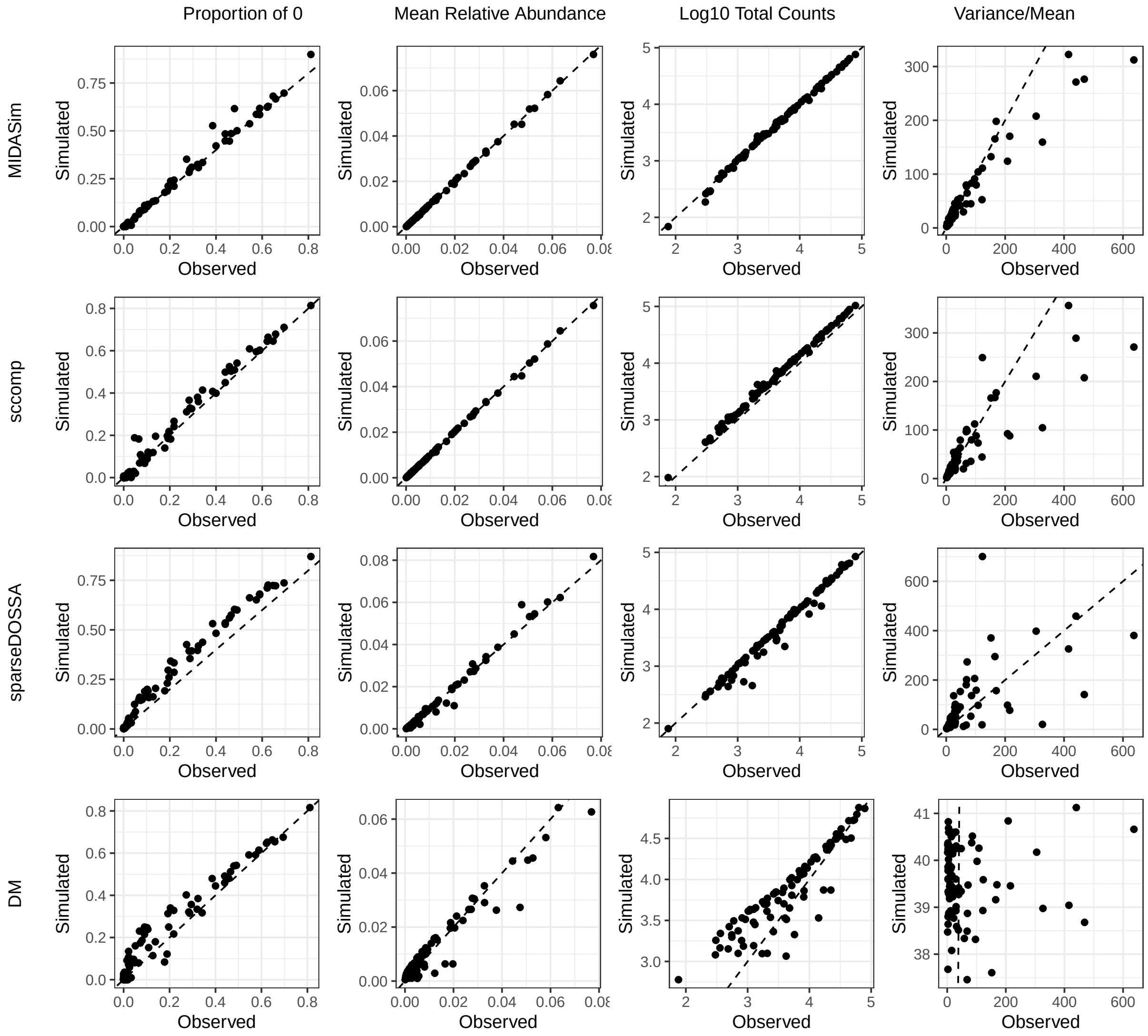
Proportion of 0
Mean Relative Abundance
Log10 Total Counts
Variance/Mean
MIDASim
sccomp
sparseDOSSA
DM
Simulated
Observed

**c**

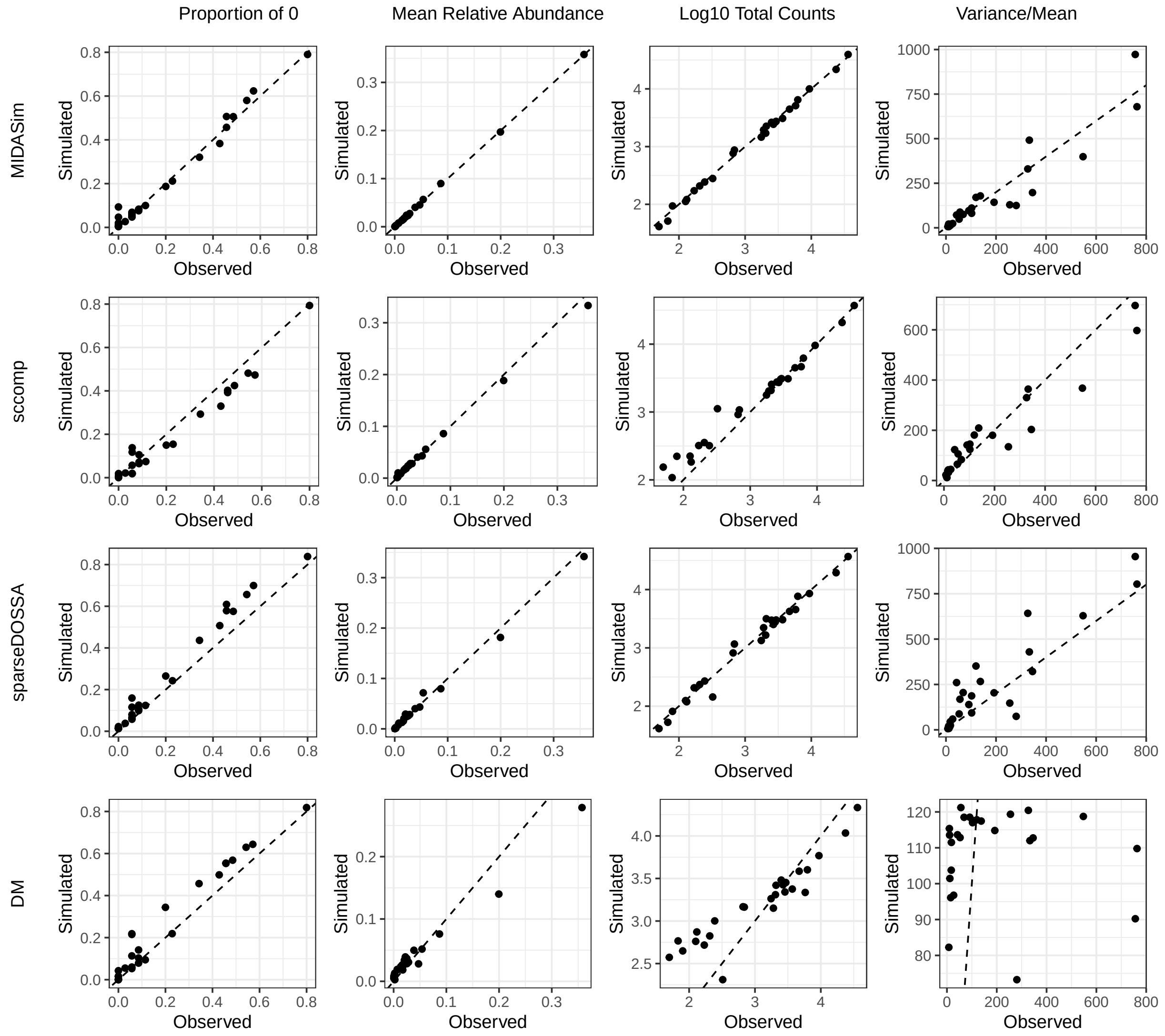
Proportion of 0
Mean Relative Abundance
Log10 Total Counts
Variance/Mean
MIDASim
sccomp
sparseDOSSA
DM
Simulated
Observed

**d**

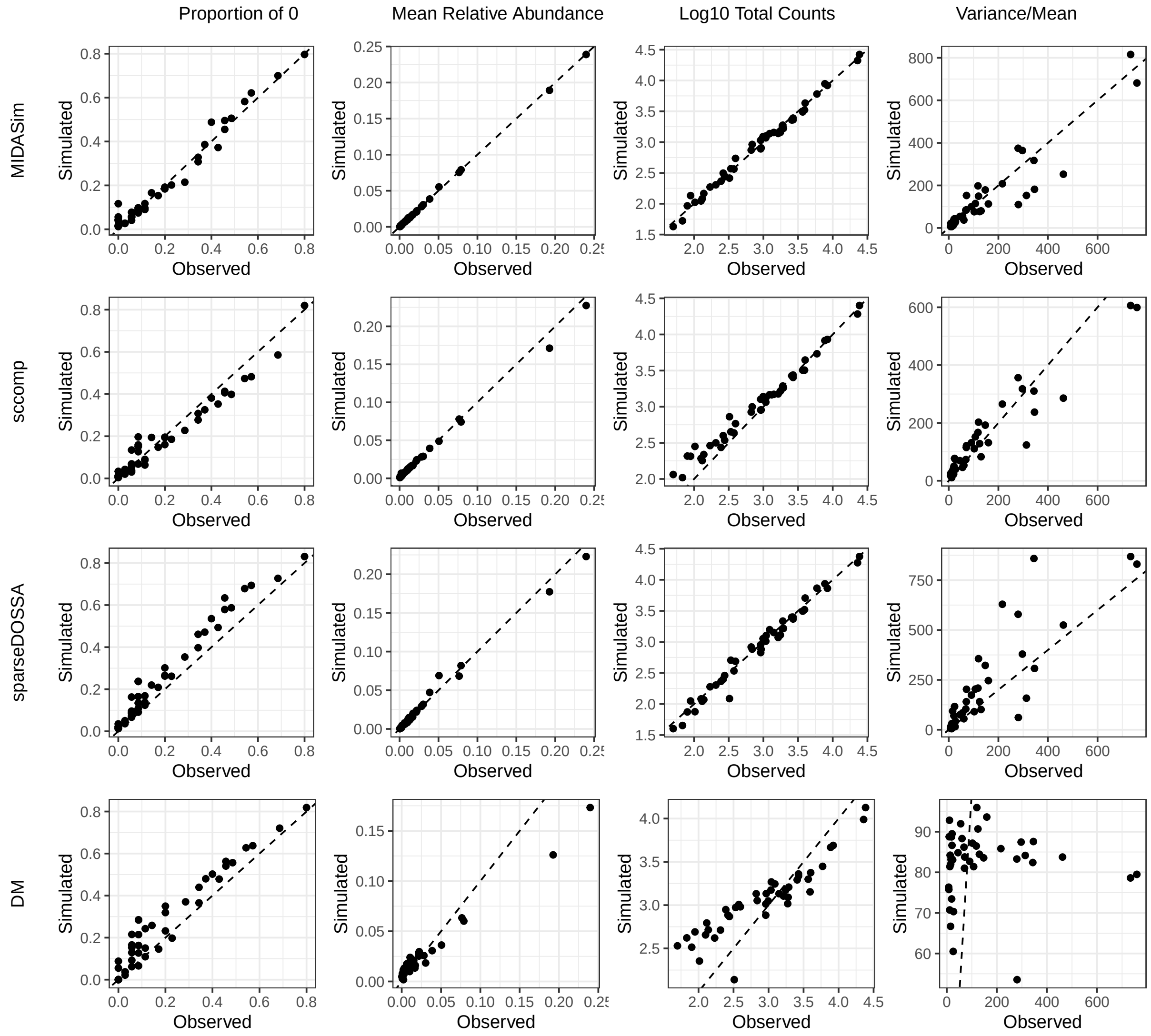
Proportion of 0
Mean Relative Abundance
Log10 Total Counts
Variance/Mean
MIDASim
sccomp
sparseDOSSA
DM
Simulated
Observed

**e**

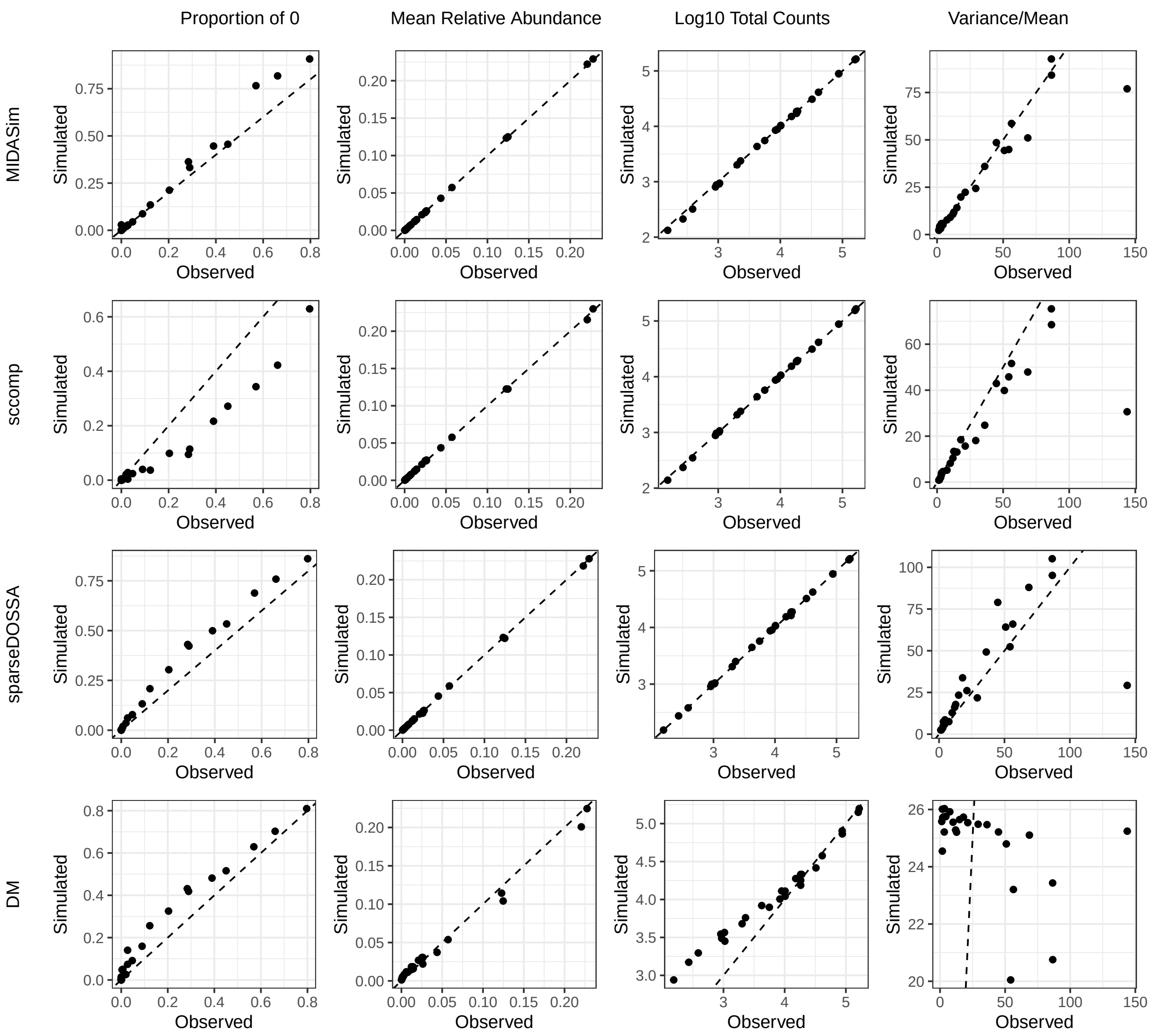


**Supplementary Fig. S8:** Comparison of cell-group features computed from observed and simulated samples using four simulators across five real datasets AD CT (**a**), AD CS (**b**), ILD CT (**c**), ILD CS (**d**), OneK1K CS **e**). Four cell-group features are shown: the proportion of zeros, mean relative abundance, $\log_{10}$ total counts, and overdispersion (variance-to-mean ratio). Each point represents a cell group; the x-axis shows estimates from observed samples, and the y-axis shows the mean estimates across 100 simulated replicates. The black dashed diagonal indicates equality between simulated and observed values.

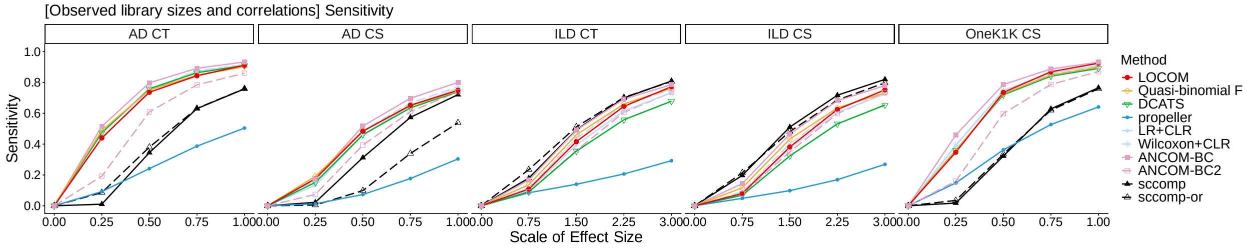


**Supplementary Fig. S9:** Sensitivity in the default setting of observed library sizes and correlations. The scale of effect size is used to scale the estimated effect sizes from real datasets to create varying strengths of signals.

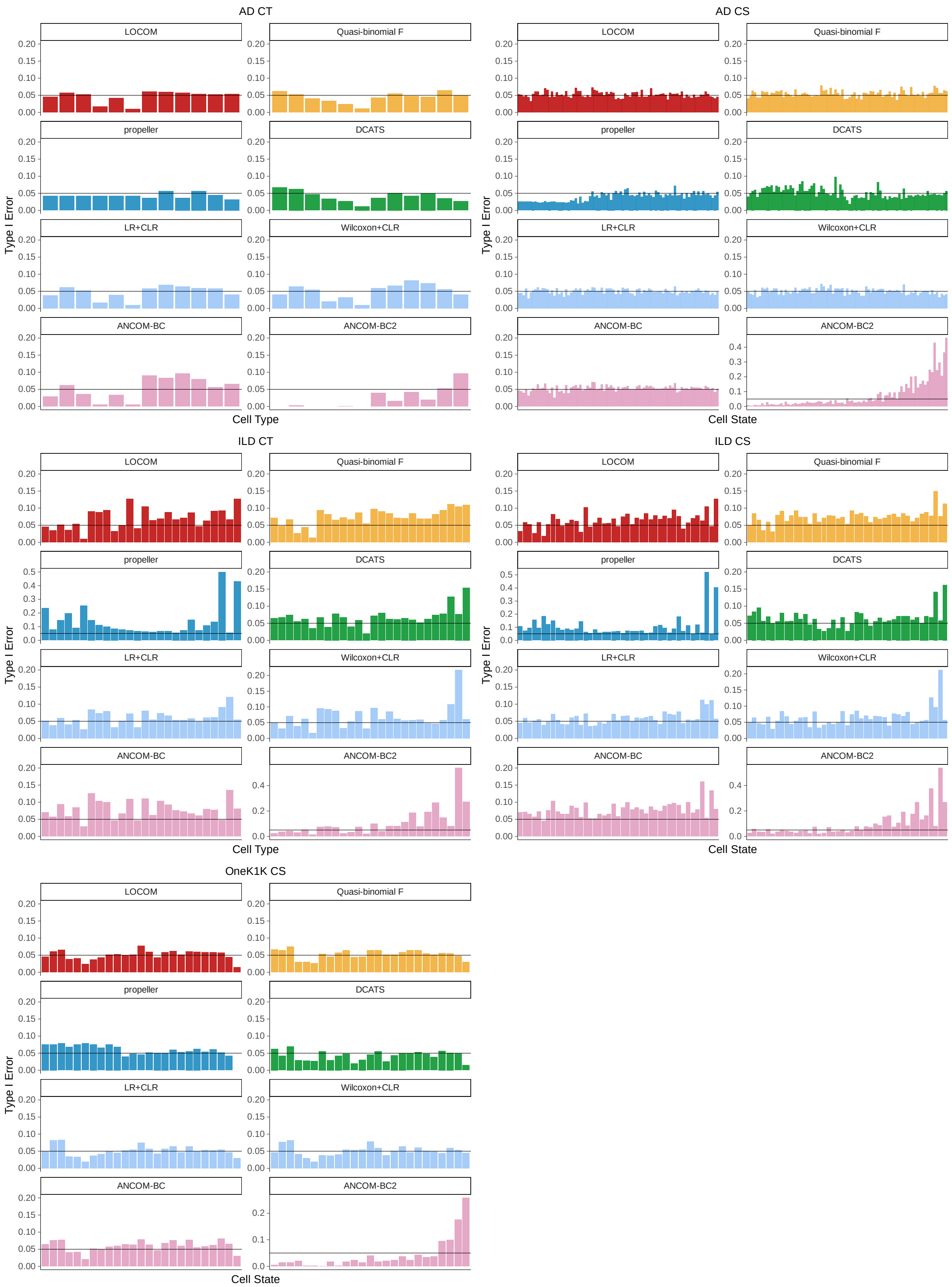


**Supplementary Fig. S10:** Type I error rates by cell groups in analysis of simulated replicates of the ILD CS, ILD CT, and OneK1K datasets under the default setting. The x-axes denote cell types and cell states in ascending order of the proportion of zero. The black lines mark the nominal 0.05 level.

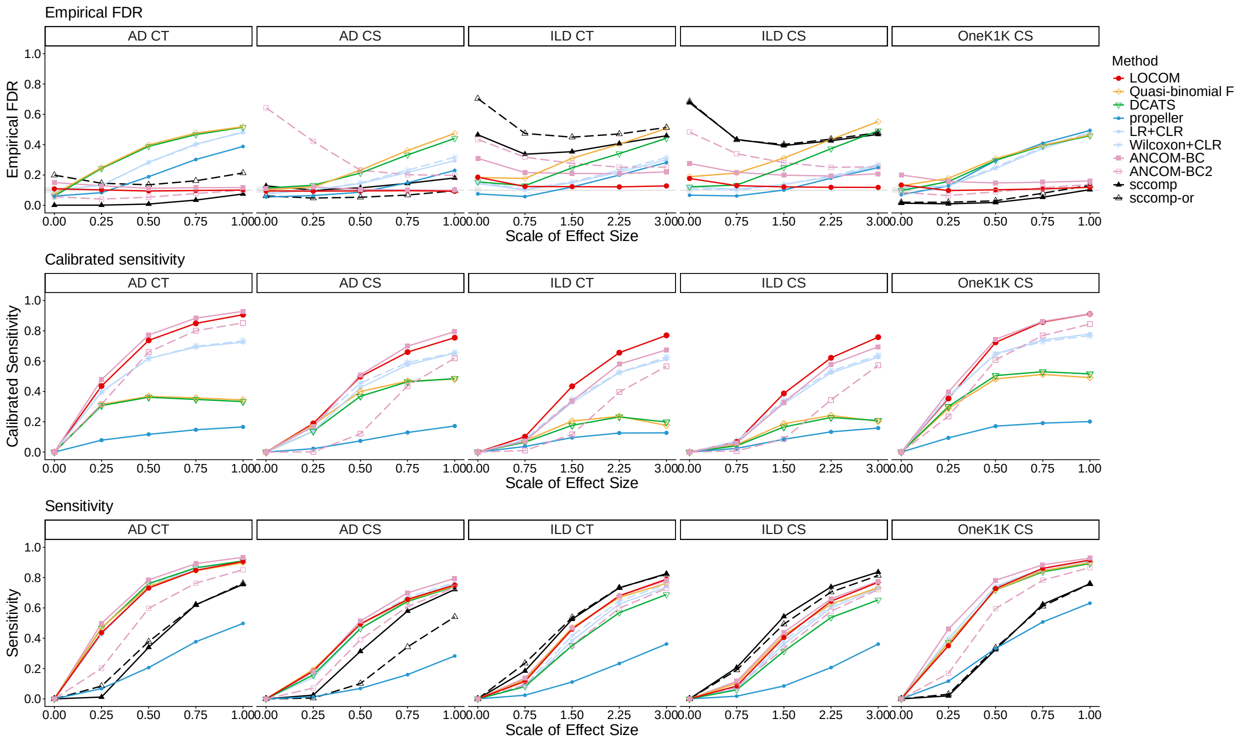


**Supplementary Fig. S11:** Empirical FDR, calibrated sensitivity, and sensitivity in analysis of simulated replicates of four real datasets in the setting with non-differential library sizes and correlations as observed in real data. Refer to the caption of **Fig. 5** for additional information.

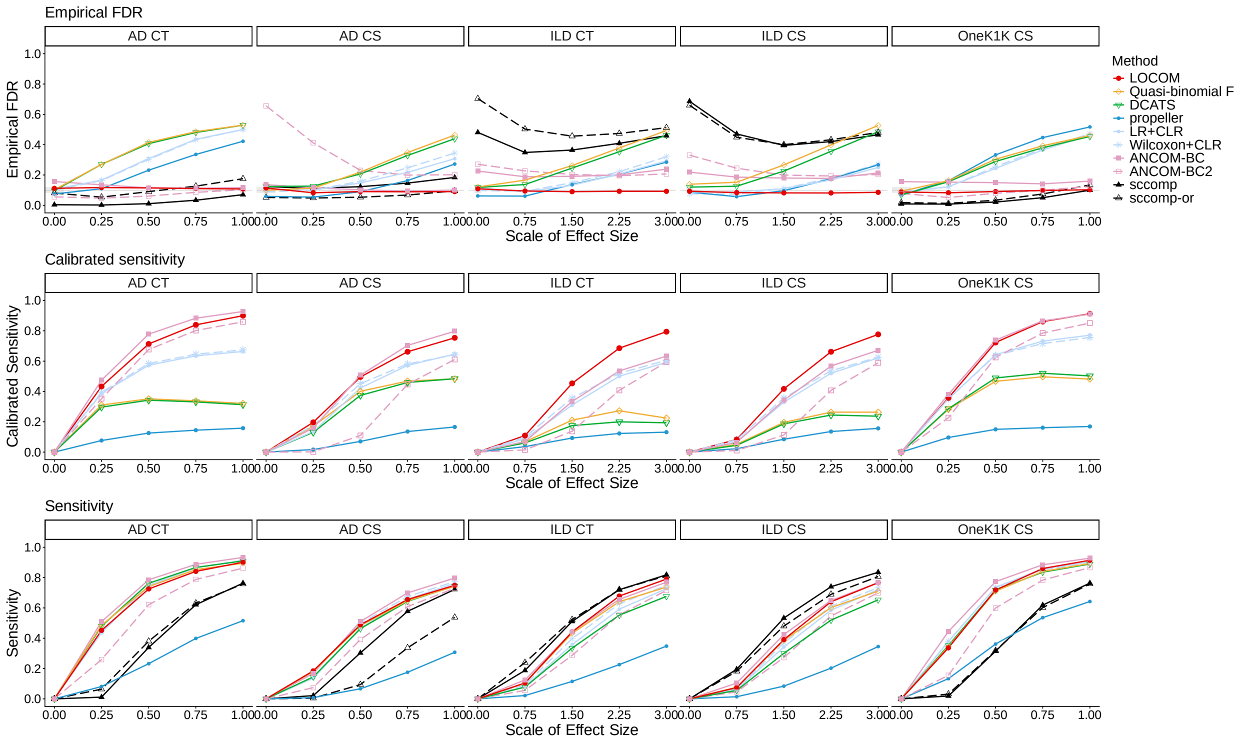


**Supplementary Fig. S12:** Empirical FDR, calibrated sensitivity, and sensitivity in analysis of simulated replicates of four real datasets in the setting with non-differential library sizes and no correlations. Refer to the caption of **Fig. 5** for additional information.

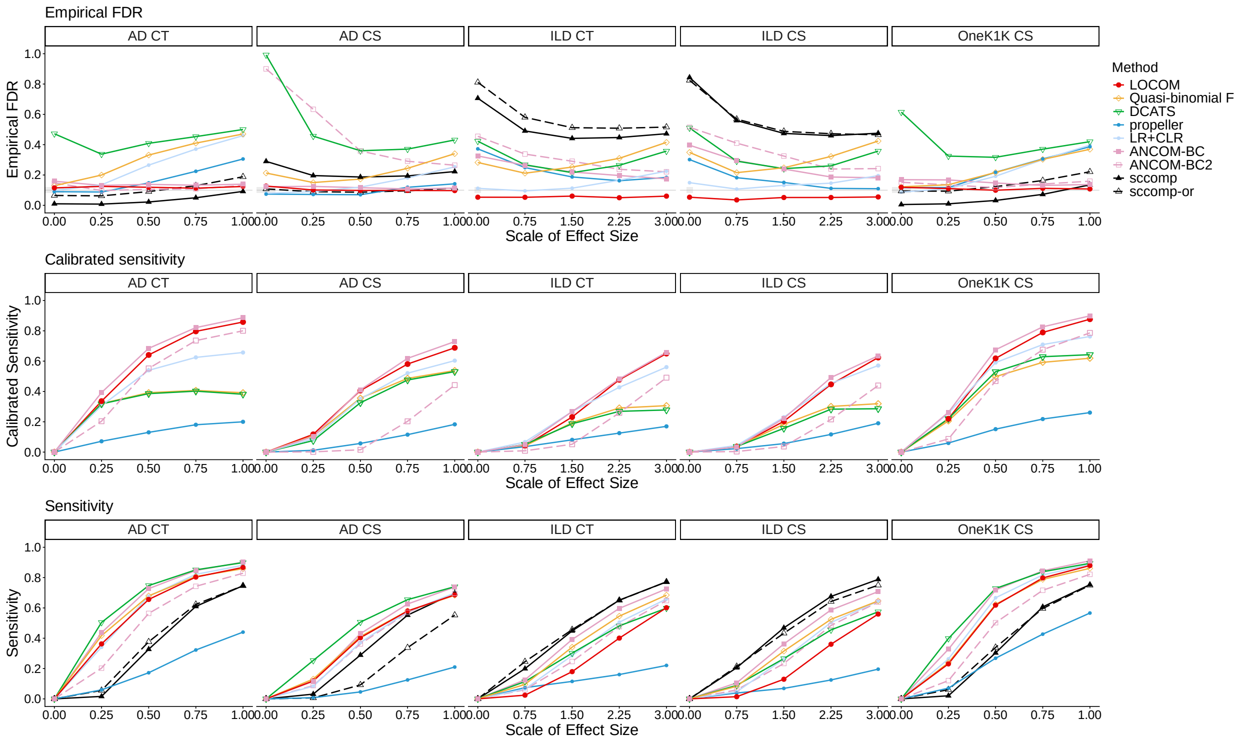


**Supplementary Fig. S13:** Empirical FDR, calibrated sensitivity, and sensitivity in analysis of simulated replicates of four real datasets in the setting with confounders. Refer to the caption of **Fig. 5** for additional information.

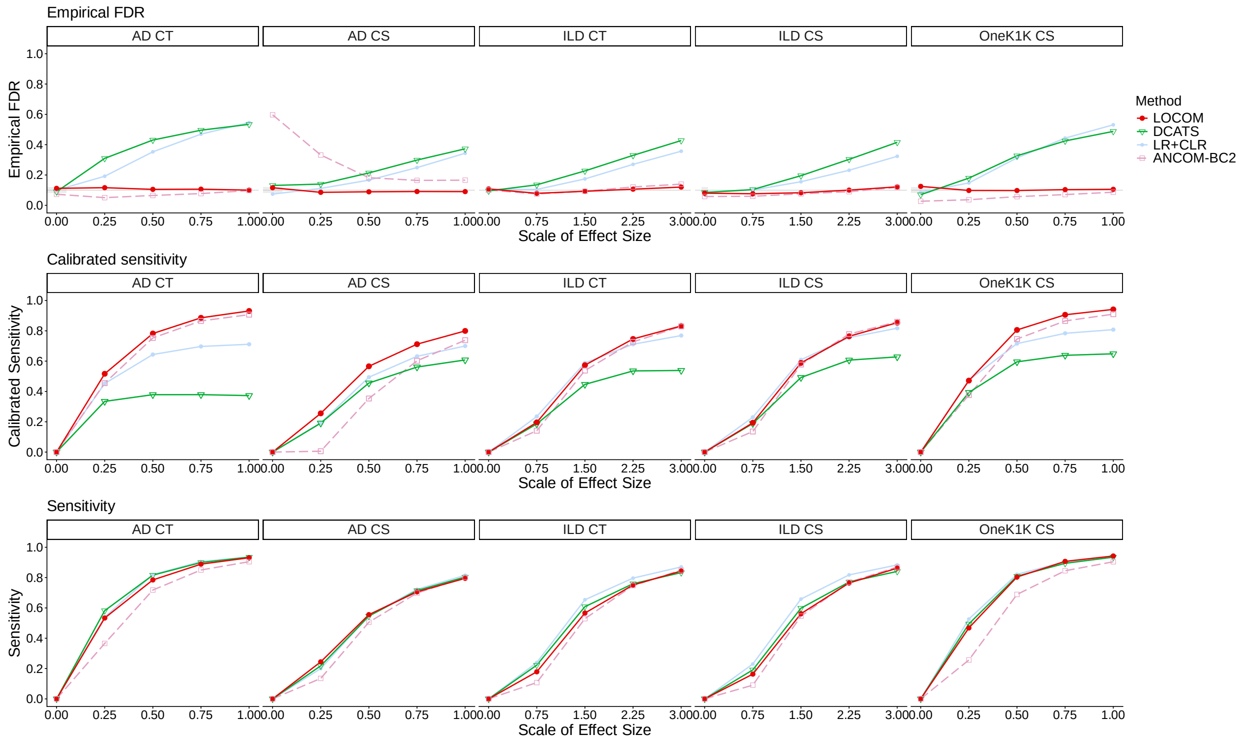


**Supplementary Fig. S14:** Empirical FDR, calibrated sensitivity, and sensitivity in analysis of simulated replicates of four real datasets in the setting with continuous traits. Refer to the caption of **Fig. 5** for additional information.

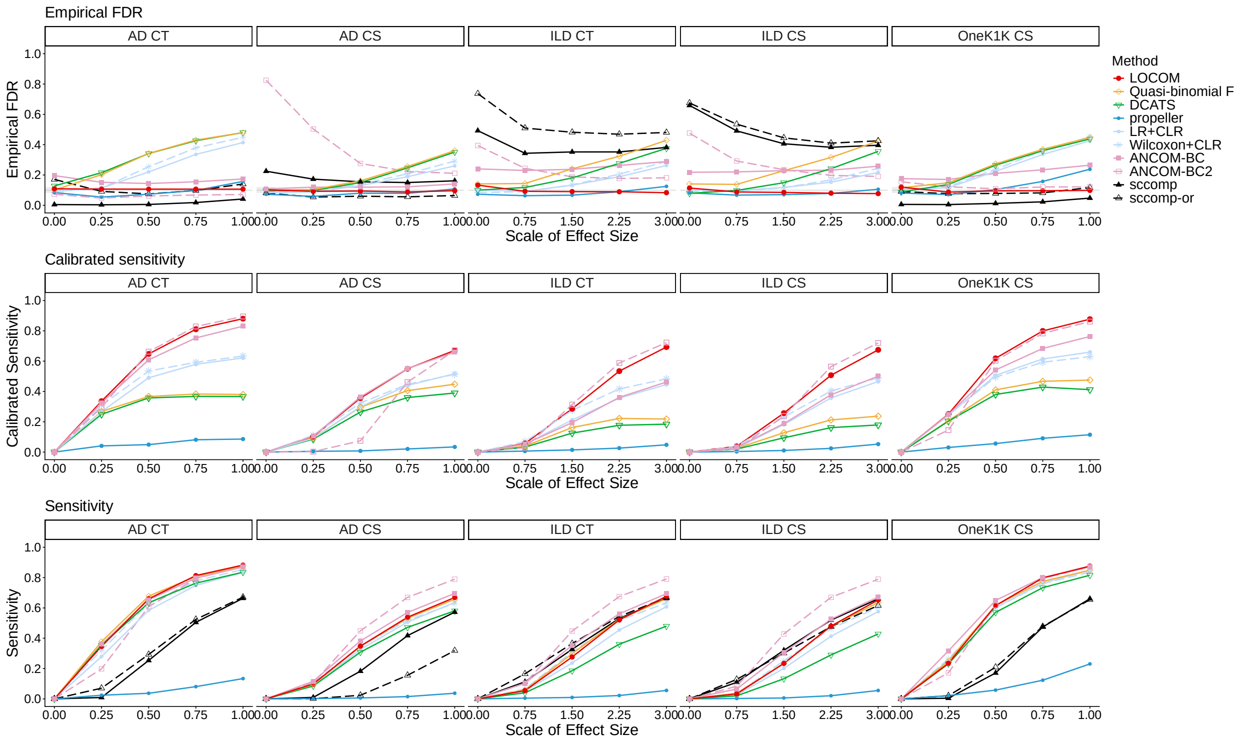


**Supplementary Fig. S15:** Empirical FDR, calibrated sensitivity, and sensitivity in analysis of simulated replicates of four real datasets in the setting with an alternative simulator (sparseDOSSA). Refer to the caption of **Fig. 5** for additional information.

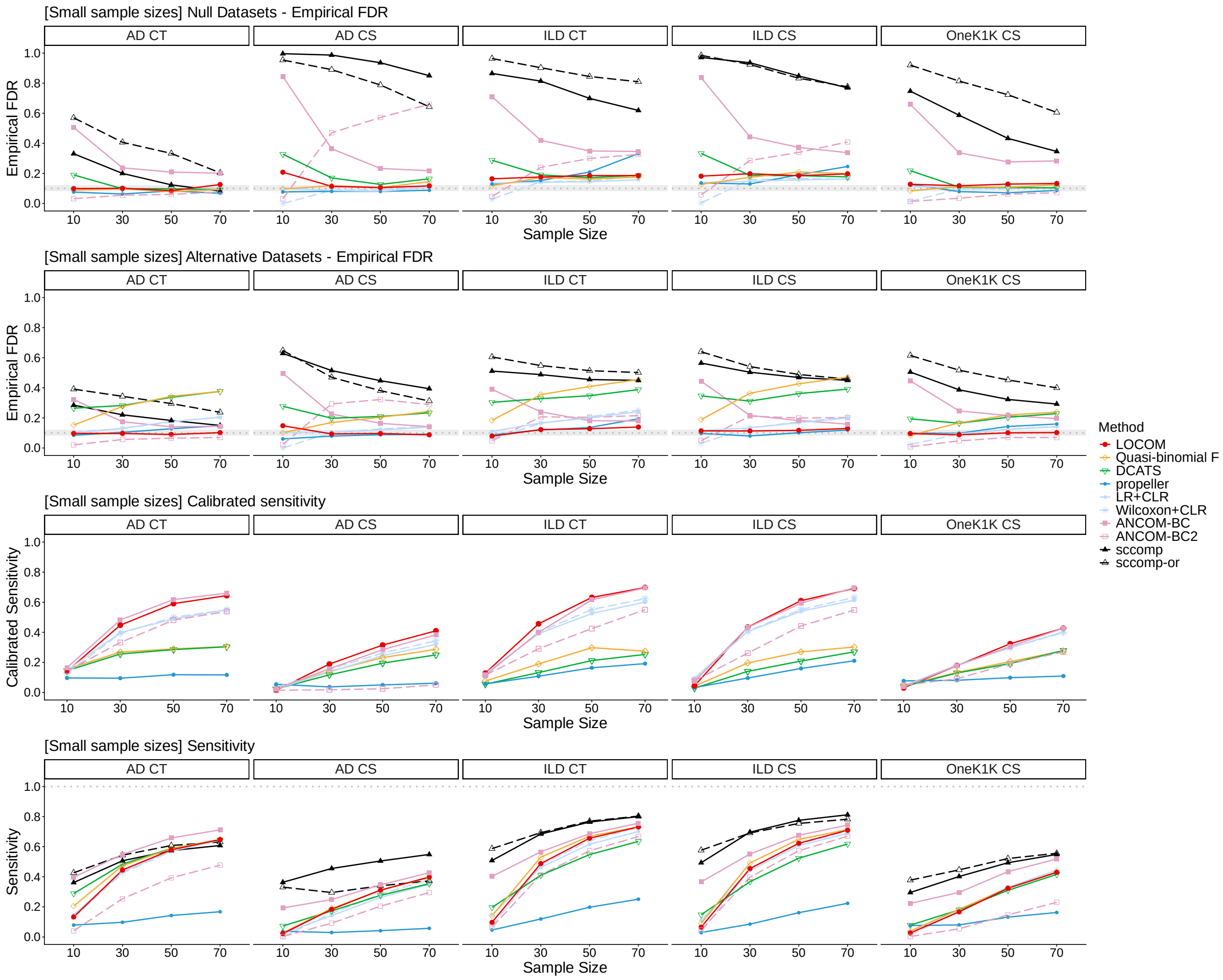


**Supplementary Fig. S16:** Empirical FDR, calibrated sensitivity, and sensitivity in analysis of simulated replicates of four real datasets in the setting with small sample sizes. The first and the second row show empirical FDR on simulated null replicates and alternative replicates, respectively. Refer to the caption of **Fig. 5** for additional information.

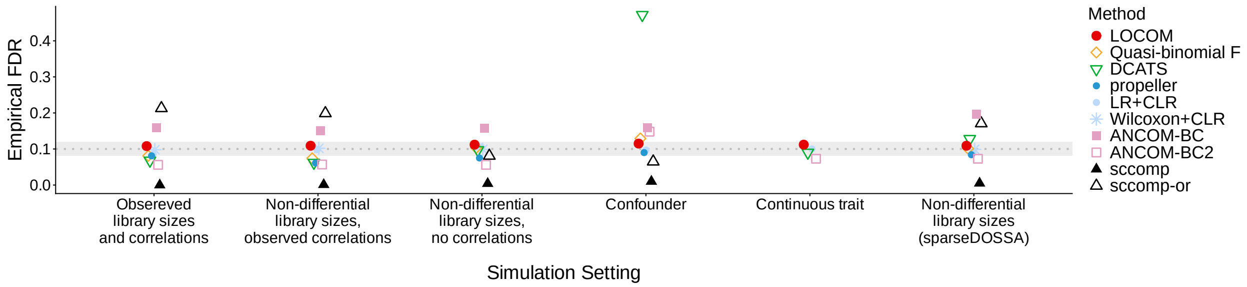


**Supplementary Fig. S17:** Empirical FDR on null datasets (scale of effect size = 0) for the AD CT scenario across different settings. The gray line represents the nominal 10% level, and the gray shaded band indicates the 95% confidence interval.

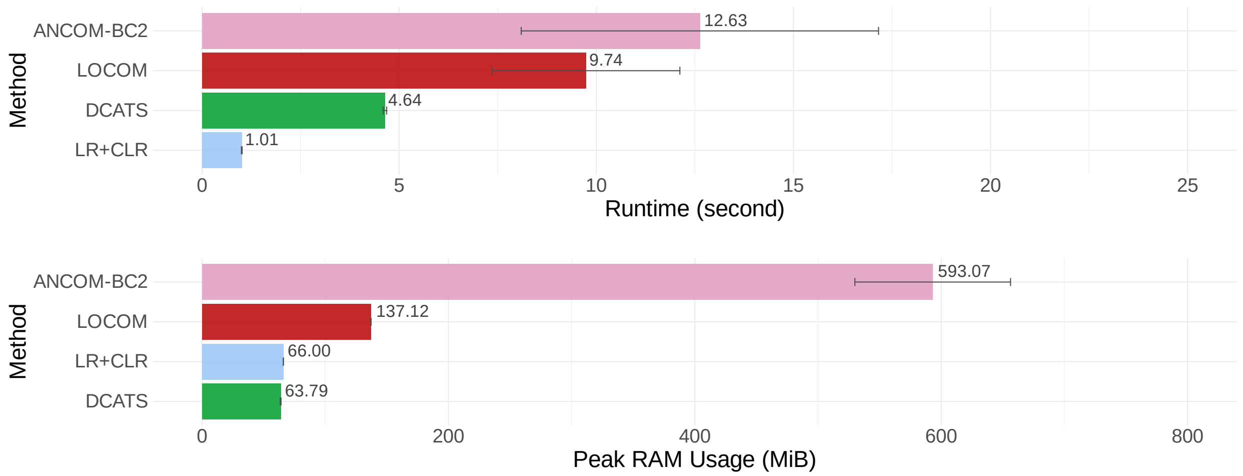


**Supplementary Fig. S18:** Runtime and memory usage in analysis of simulated alternative replicates of the AD CS dataset with a continuous trait. Refer to the caption of **Fig. 6** for additional information.

### 2 Supplementary Tables

**Supplementary Table S1:** Permutational multivariate analysis of variance (PERMANOVA).

| Template Dataset | Method | P-value |
|---|---|---|
| AD CS | MIDASim | 0.0014 |
| | sccomp | 0.0013 |
| | sparseDOSSA | 0.0010 |
| | DM | 0.0010 |
| AD CT | MIDASim | 0.8342 |
| | sccomp | 0.8369 |
| | sparseDOSSA | 0.9630 |
| | DM | 0.0010 |
| ILD CS | MIDASim | 0.7510 |
| | sccomp | 0.6171 |
| | sparseDOSSA | 0.8810 |
| | DM | 0.0028 |
| ILD CT | MIDASim | 0.7251 |
| | sccomp | 0.5167 |
| | sparseDOSSA | 0.7559 |
| | DM | 0.0097 |
| OneK1K CS | MIDASim | 0.0649 |
| | sccomp | 0.2652 |
| | sparseDOSSA | 0.9947 |
| | DM | 0.0010 |

The P-value represents the mean of 100 PERMANOVA p-values using Bray-Curtis distances across simulation replicates for each simulator and template dataset, where each simulation replicate is compared to the template dataset. DM: Dirichlet multinomial model.

**Supplementary Table S2:** Covariates included in the AD, ILD, and OneK1K real-data analyses.

| Dataset | Variable | P-value[1] |
|---|---|---|
| AD | Sex | 0.13 |
| | APOE $\epsilon 4$ | $< 0.01$ |
| | Postmortem interval | 0.40 |
| ILD | Sample source | $< 0.01$ |
| | Age | $< 0.01$ |
| OneK1K | Age | 0.60 |

[1] Detailed definitions are provided in the original dataset publications [1, 2, 3].

[2] Pearson's Chi-squared test, Wilcoxon rank sum test, and Fisher's exact test were selected as appropriate for each variable.
